# Antibiotics drive expansion of rare pathogens in a chronic infection microbiome model

**DOI:** 10.1101/2021.06.21.449018

**Authors:** John J. Varga, Conan Zhao, Jacob D. Davis, Yiqi Hao, Jennifer M. Farrell, James R. Gurney, Eberhard Voit, Sam P Brown

## Abstract

Chronic (long-lasting) infections are globally a major and rising cause of morbidity and mortality. Unlike typical acute infections, chronic infections are ecologically diverse, characterized by the presence of a polymicrobial mix of opportunistic pathogens and human-associated commensals. To address the challenge of chronic infection microbiomes, we focus on a particularly well-characterized disease, cystic fibrosis (CF), where polymicrobial lung infections persist for decades despite frequent exposure to antibiotics. Epidemiological analyses point to conflicting results on the benefits of antibiotic treatment, and are confounded by the dependency of antibiotic exposures on prior pathogen presence, limiting their ability to draw causal inferences on the relationships between antibiotic exposure and pathogen dynamics. To address this limitation, we develop a synthetic infection microbiome model, and benchmark on clinical data. We show that, in the absence of antibiotics, the microbiome structure in a synthetic sputum medium is highly repeatable and dominated by oral commensals. In contrast, challenge with physiologically relevant antibiotic doses leads to substantial community perturbation characterized by multiple alternate pathogen-dominant states and enrichment of drug-resistant species. These results provide evidence that antibiotics can drive the expansion (via competitive release) of previously rare opportunistic pathogens and offer a path towards microbiome-informed conditional treatment strategies.

## Introduction

One of the crowning achievements of modern medicine is the discovery and development of antibiotics. Our ability to control infections has opened new medical opportunities; for instance, the ability to control the risk of infection during surgery has led to greater access to safe medical interventions. Unfortunately, physicians now face two growing crises that impact their ability to treat bacterial infections. The first is widely recognized – the evolution of antibiotic resistance^1^. The second receives less attention – chronic (long-lasting) infections that are more difficult to control^2–4^.

Chronic infections are globally a rising burden on health-care systems due to increases in populations at risk (e.g., the elderly, people with diabetes or other chronic diseases)^5^. At-risk populations have deficits in host-barrier defenses and/or immune function that provide an opening for the establishment of infections, and these chronic infections are further complicated by changes in pathogen growth mode (e.g., formation of multicellular aggregates^6–8^) and development of complex multispecies communities^9^.

To address the global challenge of chronic infections, we focus on a particularly well-characterized disease, cystic fibrosis (CF), where bacterial infections can persist for decades. CF is caused by mutations in the cystic fibrosis transmembrane conductance regulator (CFTR), an ion channel that conducts chloride and thiocyanate ions across epithelial cell membranes, leading to defective mucociliary clearance and polymicrobial infection^10, 11^, resulting in eventual pulmonary failure^12, 13^.

Traditionally, CF research and patient care have focused on a small cohort of opportunistic pathogens, highlighting a distinct successional pattern^14^ characterized by peak prevalence of *Haemophilus influenzae* in childhood, *Staphylococcus aureus* during adolescence and *Pseudomonas aeruginosa* in adulthood. In addition to the core pathogen species, 16S rDNA amplicon sequencing of expectorated sputum samples has revealed much more diverse communities including numerous bacteria that are considered non-pathogenic in CF and that are normally associated with oral and upper-respiratory environments^15–19^. The functional role of these non-pathogenic taxa in CF lungs is currently disputed^20^. Epidemiological analyses have identified potentially positive roles, as higher lung function correlates with higher relative abundance of oral bacteria in sputum samples from both cross-sectional^21–23^ and longitudinal studies^24^. In contrast, *in vitro* experimental studies have suggested health risks of specific oral bacteria in the lung, due to the potential facilitation of pathogen growth^25, 26^. A third interpretation is that oral bacteria found in sputum are simply the result of sample contamination with oral microbes during expectoration^27, 28^. A number of approaches to address the sputum contamination issue have been taken, including mouth cleaning and sputum rinsing^29^, as well as more invasive sampling techniques (subject to clinical need^28, 30–32^). Most recently, computational analysis of paired sputum and saliva samples from adults with established CF lung disease has demonstrated that saliva contamination during sample collection has a minimal quantitative impact on the community profile^33^.

As a result of long-term bacterial infection, people with CF are exposed to high levels of antibiotics^34^, both as maintenance therapy^35^ and as treatment for exacerbations. In the context of a critical health challenge (an acute pulmonary exacerbation), health outcomes are variable – lung function can rapidly increase back to baseline values or remain at a new, lower baseline following antibiotic intervention. Unfortunately, a recent systematic review of 25 articles indicated little correlation between these variable clinical outcomes and antibiotic susceptibility test results for the target pathogen^36^. Several factors for this disconnect have been proposed, including differences in bacterial physiology^37^, non-representative infection sampling^38, 39^ and polymicrobial interactions^40^. In a microbiome context, epidemiological studies indicate variable outcomes of antibiotic treatment, ranging from minimal impact on microbiome structure^15, 41, 42^ to target pathogen declines, microbiome structural changes^43–46^ and risk of subsequent infection^47^. However, there is a fundamental confounding factor in these epidemiological studies, as antibiotic exposures are themselves dependent on the microbiome state of the patient. Specifically, the detection of pathogens within the microbiome will dictate antibiotic choice^48^.

Here we seek to overcome this confounding impact of pathogen detection through the development and benchmarking of a clinically relevant experimental infection microbiome model. Using this model we seek to address the key question of whether antibiotics act as drivers of pathogen and resistance expansion and community diversification given experimental control of antibiotic exposures. In addition, we seek to assess the functional role of commensal microbes in a polymicrobial infection context and, specifically, to ask whether orally-derived microbes suppress or facilitate pathogen growth.

While most experimental polymicrobial models of CF have focused on two species pathogen interactions^49–51^, some studies have developed up to 6-species models^52, 53^. These more complex models have demonstrated that species antibiotic susceptibility is not impacted by community context^53^, but their use of rich media (to facilitate single-species comparisons) raises the issue of relevance to the *in vivo* context of growth in sputum^54^. Our experimental approach begins with a “synthetic sputum” that recreates the biochemical and physical conditions of the sputum found in CF lungs^55, 56^. We then add defined combinations of 10 bacterial species that together account for over 85% of the observed bacterial diversity within the CF lung in a 77 person cohort^57^. Five of these species are established human pathogens (*S. aureus*, *P. aeruginosa*, *H. influenzae, Burkholderia cenocepacia*, and *Achromobacter xylosoxidans*), while the rest are oral microbes frequently found in CF lungs. Our communities are cultured anaerobically to capture oxygen-depleted conditions within mucus plugs^58–60^. We show that under our *in vitro* model infection conditions, oral bacteria form stable communities that suppress the growth of multiple pathogen species, and this competitive suppression is reduced by controlled antibiotic exposures, leading to multiple alternate pathogen-dominant outcomes and the non-evolutionary enrichment of antibiotic resistance.

## Results

### In the absence of antibiotics, commensal anaerobes dominate over CF pathogens

Experiments performed in the absence of antibiotics demonstrated a consistent and reproducible community structure, characterized by population expansion during the initial 48 h and a composition primarily consisting of *P. melaninogenica*, *H. influenzae*, and *V. parvula* (Figure 1). At 48 h, the total bacterial density averaged about ∼7.7×10^6^ CFU/ml (+/- 2.0×10^6^ SD), which falls within the broad range of reported bacterial densities in sputum in clinical studies (typically between 10^4^ and 10^9^ CFU/ml^57, 61, 62^). From passage 2 onwards, each replicate showed a high degree of stability through time, both in terms of total abundance and relative composition. Across replicates, we also see a striking convergence in microbiome structure. To assess consistency across the 5 replicates, we calculated coefficients of variation (CV = 100% x standard deviation / mean) for each species’ total abundance, all showing under-dispersion (i.e. standard deviation less than the mean, with an average species CV of 46% at end of the experiment, see Figure S1), consistent with stabilizing ecological forces limiting variation in species densities across replicates.

**Figure 1.**
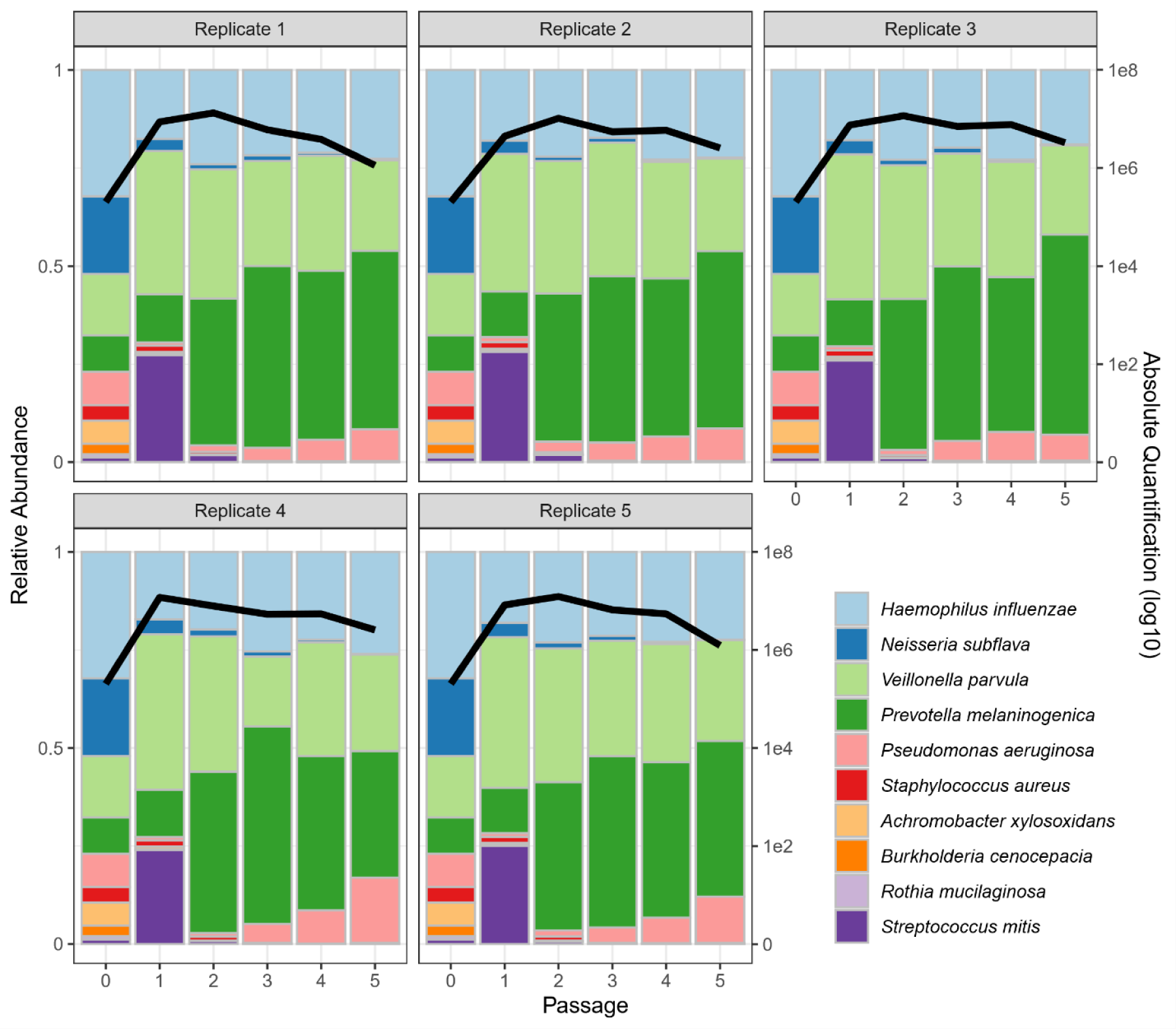
Five-fold replicated synthetic CF microbiomes converge toward a single stable state in the absence of antibiotic perturbations. 5 replicate synthetic microbiomes were grown anaerobically in artificial sputum medium. The community composition was estimated by 16S rDNA amplicon sequencing at time zero and at every two-day passage (x-axes) into fresh medium (10% transfer of 2 ml culture volume). The colored bars represent relative abundance of each species in the community (left y-axis), while the black line represents the total bacterial abundance per mL (right y-axis, log scale). Each panel represents a separate replicate experiment. Strain information is provided in Table 1 (our default *P. aeruginosa* strain is mucoid PDO300).

**Table 1:** Experimental model organisms used in synthetic community experiments. * indicates pulmonary source, ^x^ indicates oral source. Red font indicates established CF pathogen^14^. ^#^ Isolate from Children’s Hospital of Atlanta. Collectively, these organisms represent over 85% of clinical sequence reads across a 77-person CF lung microbiome study^57^.

| *Species* | *Experimental Strain* | Relative abundance of the genus in clinical samples (%) |
| --- | --- | --- |
| *Pseudomonas aeruginosa* | PDO300 (mucoid)<br>PAO1 (wildtype) | 29.7 |
| *Veillonella parvula* | Clinical<sup>#</sup> | 9.8 |
| *Rothia mucilaginosa* | ATCC49042<sup>*</sup> | 9.1 |
| *Prevotella melaninogenica* | ATCC25845<sup>*</sup> | 8.4 |
| *Streptococcus mitis* | ATCC49456<sup>x</sup> | 7.9 |
| *Haemophilus influenza* | ATCC10211 | 5.8 |
| *Staphylococcus aureus* | SAJE2 | 5.6 |
| *Achromobacter xylosoxidans* | ATCC27061 | 4.8 |
| *Neisseria subflava* | ATCC49275<sup>x</sup> | 1.8 |
| *Burkholderia cenocepacia* | K56-2 | 1.1 |

The results in Figure 1 point to a robust community structure in the absence of perturbations, consistent with the frequent dominance of oral bacteria in individuals with higher lung function but far from capturing the diversity of microbiome structures observed across the broader CF community^22, 24, 32, 57, 63^. To assess the role of variable pathogen strain identity or presence / absence, we repeated the experiments in Figure 1 with wildtype (non-mucoid) PAO1, and also with variation in the presence or absence of *S. aureus*. These manipulations produced small quantitative variations in community structure, but we observed overall the same qualitative pattern with consistent dominance by *H. influenzae*, *P. melaninogenica*, and *V. parvula* (Figure 2, Figure S2).

**Figure 2.**
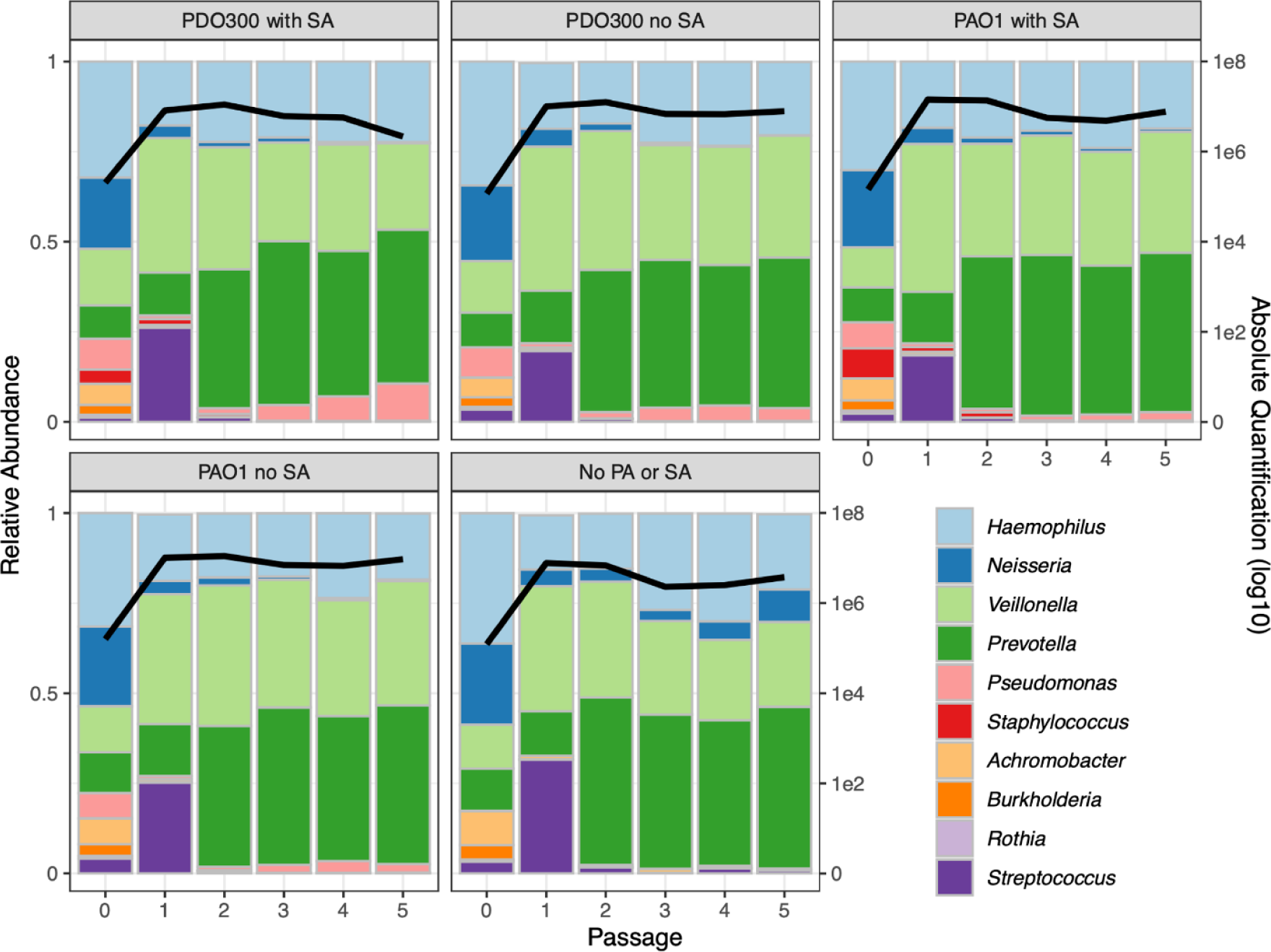
Varying the pathogen composition has minimal impact on community composition. Each panel represents the average of 5 replicates in the absence of antibiotics, the PDO300 with SA panel is the average of Figure 1. Figure details are the same as described for Figure 1. Data on individual replicates per treatment are presented in Figure S2. PAO1 and PDO300 = *P. aeruginosa* strains PAO1 and PDO300, respectively. SA = *S. aureus*.

### Antibiotics skew community structure toward pathogen expansion and dominance

Having established the repeatability and stability of the community in the absence of antibiotics, we then assessed the impact of antibiotic treatment on community structure. Communities were challenged with 3 individual antibiotics and 2 pairs commonly used in the CF clinic (tobramycin, meropenem, ciprofloxacin, tobramycin and meropenem, tobramycin and ciprofloxacin)^34, 64^ in physiologically relevant concentrations^65–67^. These perturbations resulted in dramatically different outcomes compared to the antibiotic free communities (Figure 3). Looking first at total bacterial abundance, treatments including meropenem caused the most decreases, with densities dropping to 1×10^3^ bacteria/ml at 3^rd^ passage before recovering to above 10^4^ at the final time point (Figure 3, black lines).

**Figure 3.**
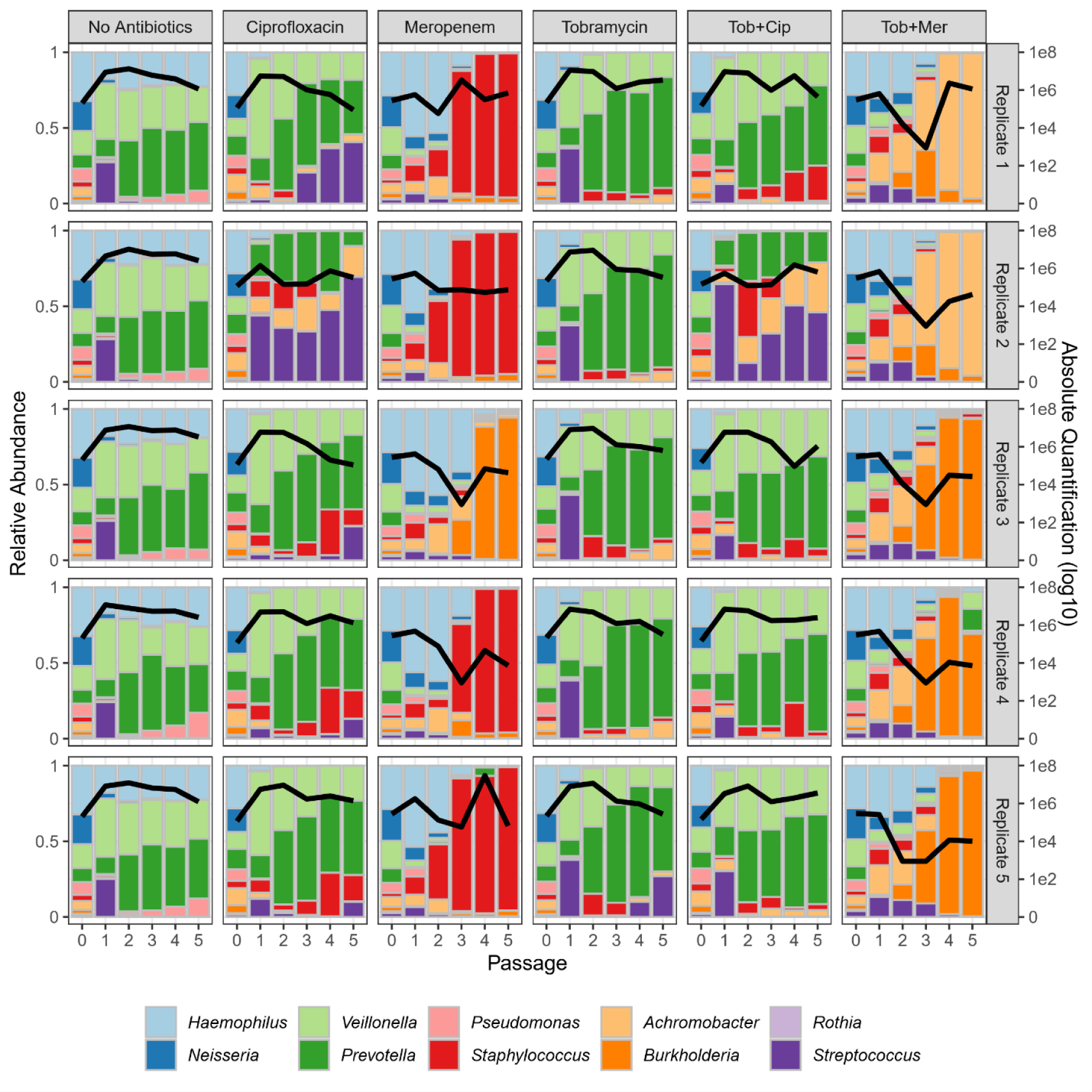
Antibiotic treatments produce large community fluctuations and alternative community states. Columns represent distinct antibiotic treatments (the first ‘no antibiotics’ control column is reproduced from Figure 1), rows represent 5 replicates. The left axes measure community composition (bar charts), the right axes measure total bacterial abundance per mL (black lines). Experimental procedures, sampling, and analysis were performed as described in Figure 1. Fresh antibiotics were re-supplemented at each passage. Total abundance data by species is presented for each treatment and timepoint in Figure S3.

Turning next to compositional structure, we see not only that the responses to different antibiotics are different but that the same antibiotic treatment often leads to distinct end points across replicates. For instance, experimental treatment with meropenem plus tobramycin leads to dominance by either *B. cenocepacia* (3 replicates) or *A. xylosoxidans* (2 replicates). Both outcomes are very concerning for a patient^68–70^.

Figure 3 indicates large shifts in response to antibiotic treatments, but compositional analysis alone cannot separate the relative importance of differential survival versus differential expansion. To address this distinction, we examined species absolute abundance data (Figure S4).

Using absolute abundances, we can now test whether pathogens undergo competitive release (expansion, following removal of competitors^71–73^) in response to antibiotic exposure, by assessing whether the final pathogen density is greater in the presence of antibiotic compared to its absence (Figure 4). Comparing mean densities (across replicates), we find evidence for significant competitive release of *S. aureus* when exposed to tobramycin plus ciprofloxacin (on average a 36-fold increase compared to no-antibiotic control). For *B. cenocepacia,* we see competitive release under the tobramycin plus meropenem treatment (19-fold increase), and for *A. xylosoxidans*, we see competitive release under tobramycin (25-fold increase) and tobramycin plus ciprofloxacin treatments (35-fold increase). We also see competitive release of *S. mitis* in ciprofloxacin (Figure S4). In contrast, there is evidence of significant suppression of *H. influenzae* and *P. aeruginosa* in all antibiotic treatments (Figure 4; two-tailed Wilcoxon test, *p* < 0.01), together with *N. subflava* in all treatments as well as *V. parvula* and *P. melaninogenica* in all meropenem treatments (Figure S4).

**Figure 4.**
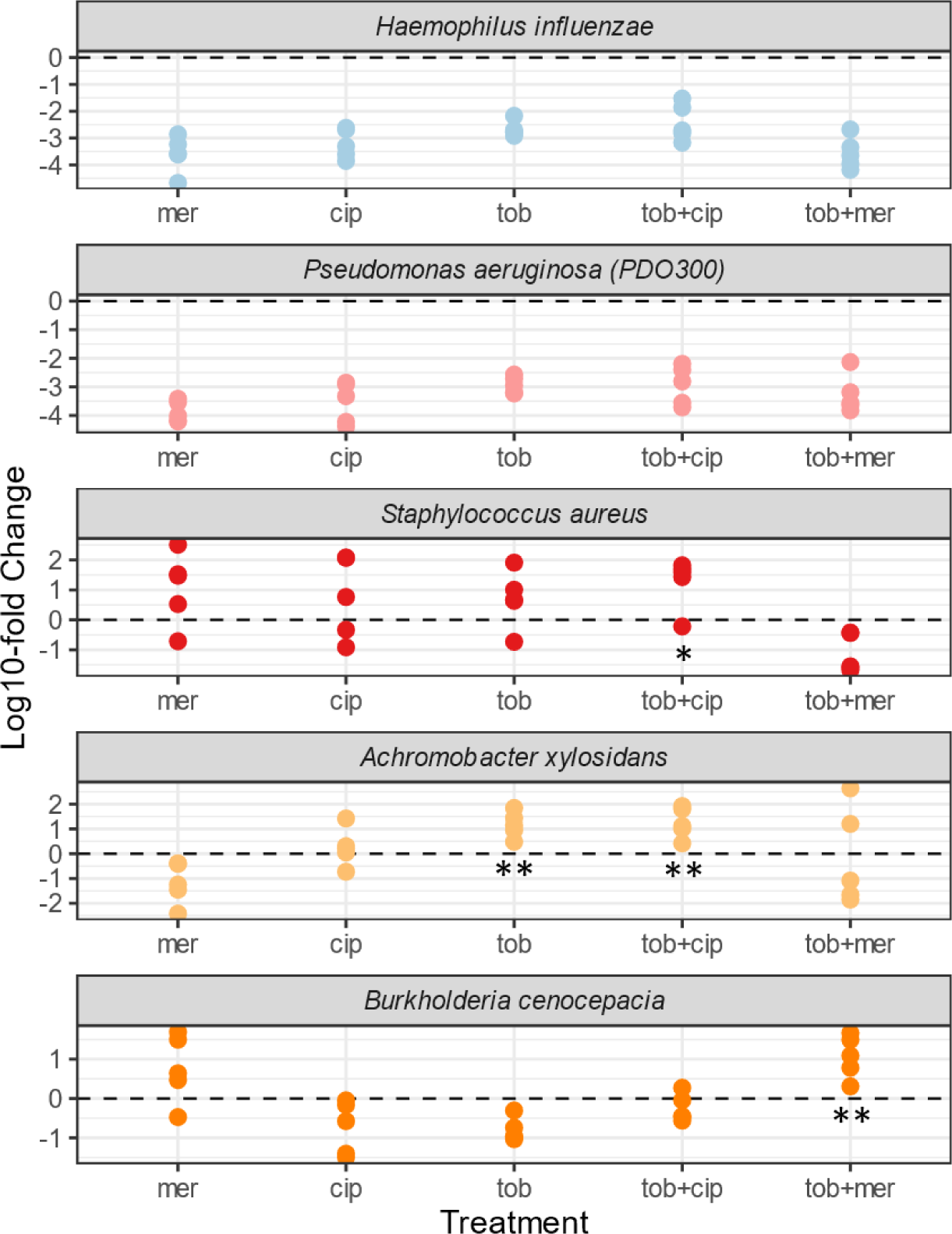
Absolute pathogen densities are often increased under antibiotic exposures. Each dot corresponds to the fold-change difference of an individual replicate of species-specific final time-point absolute density under defined antibiotic treatments, compared to the mean value of the no antibiotic control (data redrawn from Figure 3). Mer = meropenem, cip = ciprofloxacin, tob = tobramycin. Asterisks denote significantly higher final densities in presence of antibiotic, compared to antibiotic-free controls (competitive release; one-tailed Wilcoxon test * *p* < 0.05, ** *p* < 0.01).

In addition to changes in mean species densities in response to antibiotic treatments, we can also see large variability across replicates (Figures 3, 4). In the absence of antibiotics, final absolute densities are tightly clustered within an order of magnitude (average taxon CV = 46.4%, Figures S1, S4). Under antibiotic exposure, we find variation across multiple orders of magnitude (Figure S4) and over-dispersed taxon CVs (from 102% in the ciprofloxacin treatment to 135% in the tobramycin plus meropenem treatment, Figure S1).

These large fluctuations in absolute abundance lead to substantial shifts in compositional structure across replicates (Figure 3). For example, under meropenem 4 out of 5 replicates result in persistent *S. aureus* dominance, while one replicate shows persistent *B. cenocepacia* dominance. A similar signature of alternative community states is present under the tobramycin plus meropenem treatment, with trajectories split between *B. cenocepacia* and *A. xylosoxidans* dominance. To compare antibiotic treatments versus our antibiotic-free controls at a community scale, we calculated the ANOSIM R statistic^74–76^, which represents a ratio of within-treatment differences to between-treatment differences. Consistent with multiple significant impacts on the species scale (Figures 4, S4) we find significant differences between antibiotic-free and antibiotic-exposed community structures at the community scale, with treatments involving meropenem producing the largest effects (Figure S5).

### Antibiotic susceptibility partially explains community composition

The simplest hypothesis to account for the substantial impacts of antibiotic exposures on community structure (Figures 2-4) is that antibiotics present a survival filter through which only resistant organisms can pass. Under this model, the community structure after antibiotic treatment is simply the product of whether or not each taxon can grow in the antibiotic(s) administered.

To assess the survival filter hypothesis, we derived antibiotic susceptibility measures (minimal inhibitory concentrations, MICs) under standard growth conditions that allowed the more fastidious strains to grow independently (Table S2) and used these data to predict experimental responses to defined antibiotic exposures (Figure 5).

**Figure 5.**
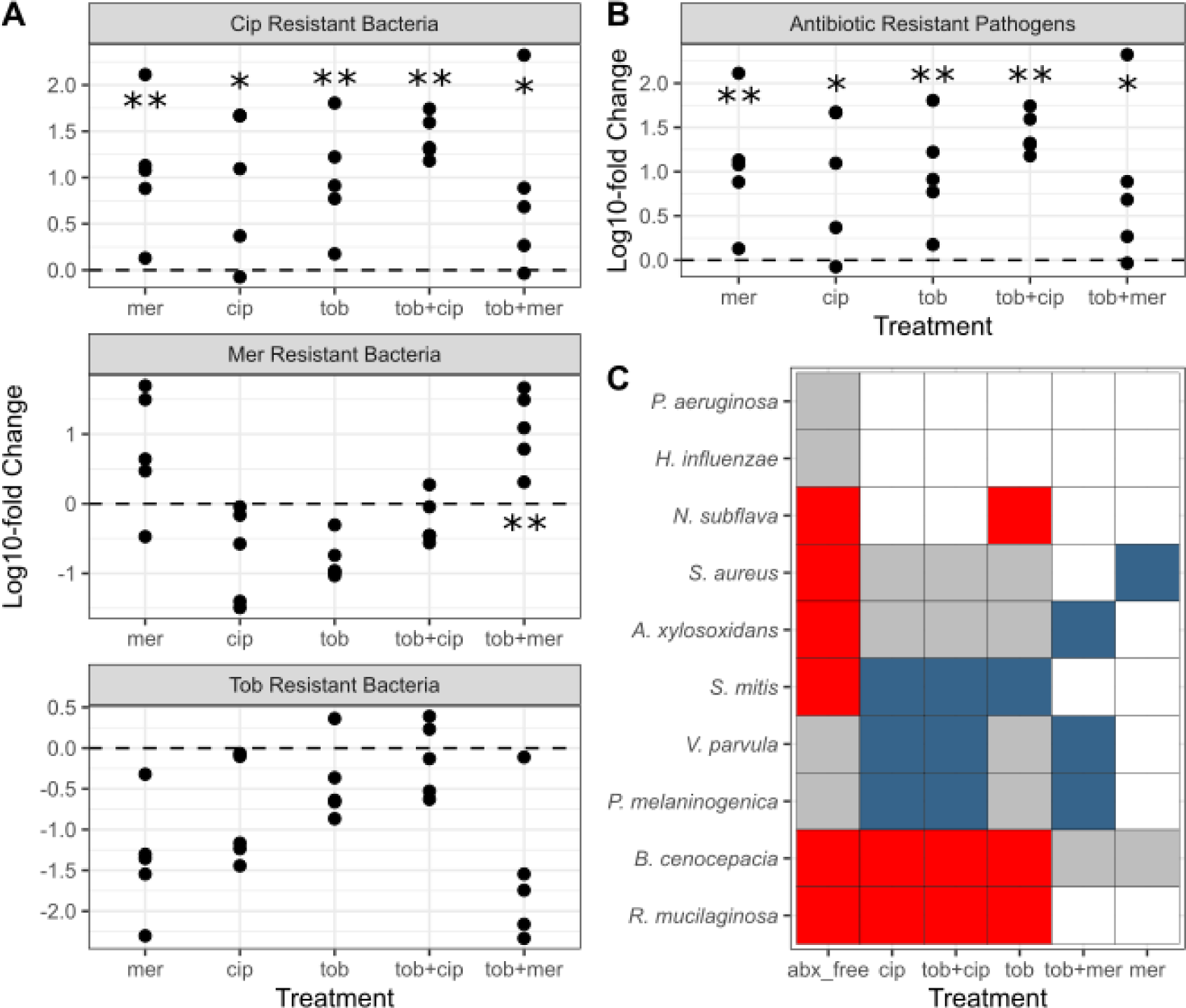
Antibiotic resistance testing partially predicts responses to antibiotic exposures. (A) Fold-change differences for specific resistance classes, compared to no antibiotic control, details as in Figure 4. Ciprofloxacin resistant bacteria are organisms with an MIC (Table S2) greater than the cip treatment level, namely *B. cenocepacia, A. xylosoxidans, S aureus, and R. mucilaginosa*. Taking the same approach, tob resistant organisms are *B. cenocepacia, A. xylosoxidans, S aureus, and R. mucilaginosa*, *P. melaninogenica, N. subflava,* and *V. parvula.* Mer resistant organisms are *B. cenocepacia* alone. (B) Drug-resistant pathogens are *B. cenocepacia, A. xylosoxidans,* and *S aureus.* (C) For each species / drug combination, we assessed predicted survival (MIC in rich medium (Table S2) > experimental concentration) and observed survival (relative abundance of at least 1% averaged across all five replicates at the final time point (Figure 3)). True positive cases (predicted and observed present) are coded in grey, true negatives (predicted and observed absent) in white. False positives (predicted present, observed absent – evidence for competition) are in red, and false negatives (predicted absent, observed present – evidence for facilitation) are in blue. Species order was determined through clustering via stringdist^77^.

In a first analysis, we pooled species (pathogens and commensals) by their drug resistance states, and ask whether organisms with specific drug resistances were enriched under antibiotic exposures, compared to the no antibiotic control (Figure 5A). Here we see data consistent with the survival filter for ciprofloxacin and meropenem resistant organisms, but not for tobramycin. We next pool drug resistant pathogens together (*S. aureus, B. cenocepacia*, and *A. xylosoxidans*) and find consistent enrichment (19- to 41-fold on average per treatment) across all drug exposures (Figure 5B).

We next turn to individual species, and ask whether their resistant/susceptible status can effectively predict their presence/absence under defined drug exposures (Figure 5C). Figure 5C illustrates that the lab strain PDO300 behaves as expected given its overall antibiotic susceptibility – it is present in the absence of treatment but then absent (average relative abundance is <1%) in the presence of antibiotics. The same is true for *H. influenzae*. However, for multiple examples the ability to resist antibiotics (in a standard clinical assay^78, 79^) did not account for the presence / absence of the species after treatment. In red, Figure 5C displays cases where the species was predicted to be present (given MIC resistance data, Table S2), but was nevertheless absent in the final community. This pattern is suggestive of an additional role for microbe-microbe competitive interactions in shaping community structure, and was observed for 6 of the 10 taxa, and most often in the absence of antibiotics. Conversely, blue regions in Figure 5C identify cases where the pathogen was predicted from MIC data to be unable to grow in the allocated antibiotic, and yet was present in the multi-species community experiment in at least 1 community. This pattern is indicative of community-dependent faciliatory interactions, where other species aid the focal species to survive under antibiotic insult, for example via antibiotic detoxification^80–85^. The explanatory potential of the survival filter model is further weakened by the high variability and multiple stabile end points across replicates in response to antibiotics (Figures 3, 4), indicative of community-mediated processes^86, 87^. In the discussion we explore potential contributing reasons other than community ecological interactions for the disconnect between MIC predictions and observed community presences.

### Community compositions across all antibiotic treatments are consistent with diversity across clinically observed *in vivo* communities

We finally ask, how do our *in vitro* synthetic microbiomes compare with the diversity of microbiome structures observed in people with CF? We begin with a PCA ordination plot to visualize experimental data (initial and final timepoints from Figure 3) alongside clinical data (Figure 6).

**Figure 6.**
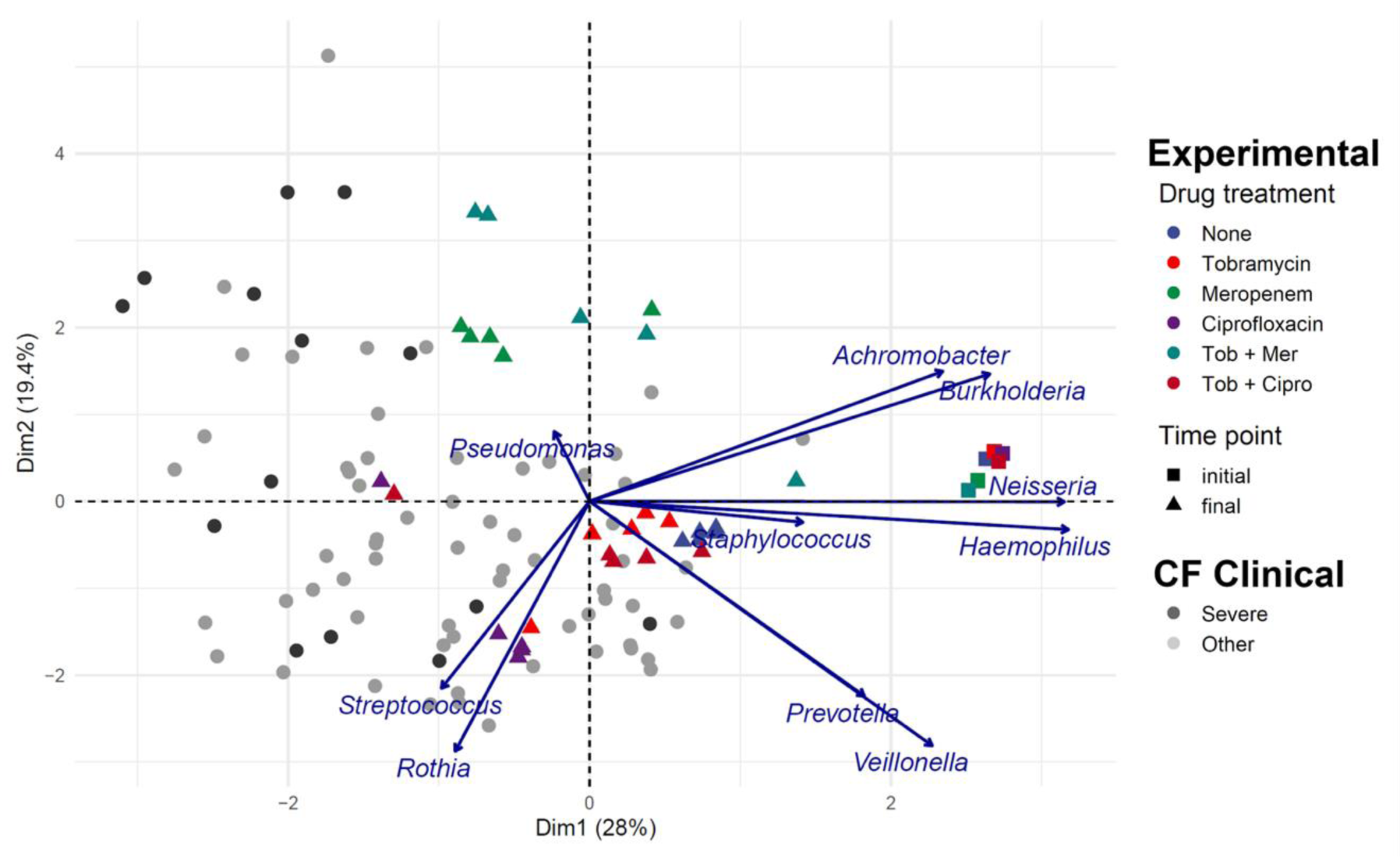
Antibiotics drive pathogen enrichment in experimental microbiomes, producing community structures that overlap with clinical sputum communities. PCA visualization of experimental microbiome data (colored triangles and squares, summarizing data in Figure 3) plus clinical microbiome data across a cohort of 77 people with CF (grey/black circles, black/severe signifies low lung function^57^). Squares illustrate experimental initial conditions, triangles are final compositions after 5 serial passages (10 days). Colors denote experimental condition (see key). Each experimental treatment is replicated 5-fold, producing highly repeatable dynamics in the absence of antibiotics (blue triangles) and variable pathogen enriched outcomes following antibiotic treatment. Antibiotics were supplemented at each passage at clinically relevant concentrations (meropenem, 15 μg / ml; tobramycin, 5 μg / ml; ciprofloxacin, 2.5 μg / ml). Each point is a single microbiome sample (species resolution for clinical samples via the DADA2 plugin in QIIME 2^57, 88^). Ordination is PCA of centered log-ratio transformed relative abundances.

Figure 6 illustrates that the ensemble of final timepoint experimental data (colored triangles; including variable responses to antibiotic perturbations) broadly overlap with clinical data. To assess differences among community structures, we again use the ANOSIM metric of community similarity R^75^, which ranges from -1 (all differences are within-groups) to 1 (all differences are between-groups). Contrasting clinical versus pooled experimental data we see a significant difference of R = 0.28, which is intermediate between the small impact of pathogen species manipulations (R values of 0.07 to 0.19 for Figure 2 data) and the large impact antibiotic manipulations (R values of 0.42 to 0.75 for Figure 3 data). For a summary of ANOSIM R statistics, see Figure S5.

Building on the overview provided by Figure 6, we now look at a more granular level and ask for each taxon whether the relative abundances (across all treatments) in our experimental model fall within the range of clinical variation from our previous clinical study^57^. We find that for 7 out of 10 taxa, our experimental model produces ranges of relative abundances that do not significantly differ from clinical data (Figure 7), thus supporting our experimental model for these taxa. In contrast, we find that our model significantly over-represents *Prevotella* and under-represents *Pseudomonas* and *Rothia.* These misses provide an opportunity to improve our model in future work, by pointing towards an environmental mismatch on oxygenation (with the strict anaerobe *P. melaninogenica* benefitting and the facultative anaerobes *P. aeruginosa* and *R. mucilaginosa* suffering from the anaerobic atmosphere). This pattern of misses suggests that the distribution of oxygenation experienced clinically by CF microbiomes is more oxygenated than that provided by our anaerobic chambers with only brief exposures to oxygen every 48 h.

**Figure 7.**
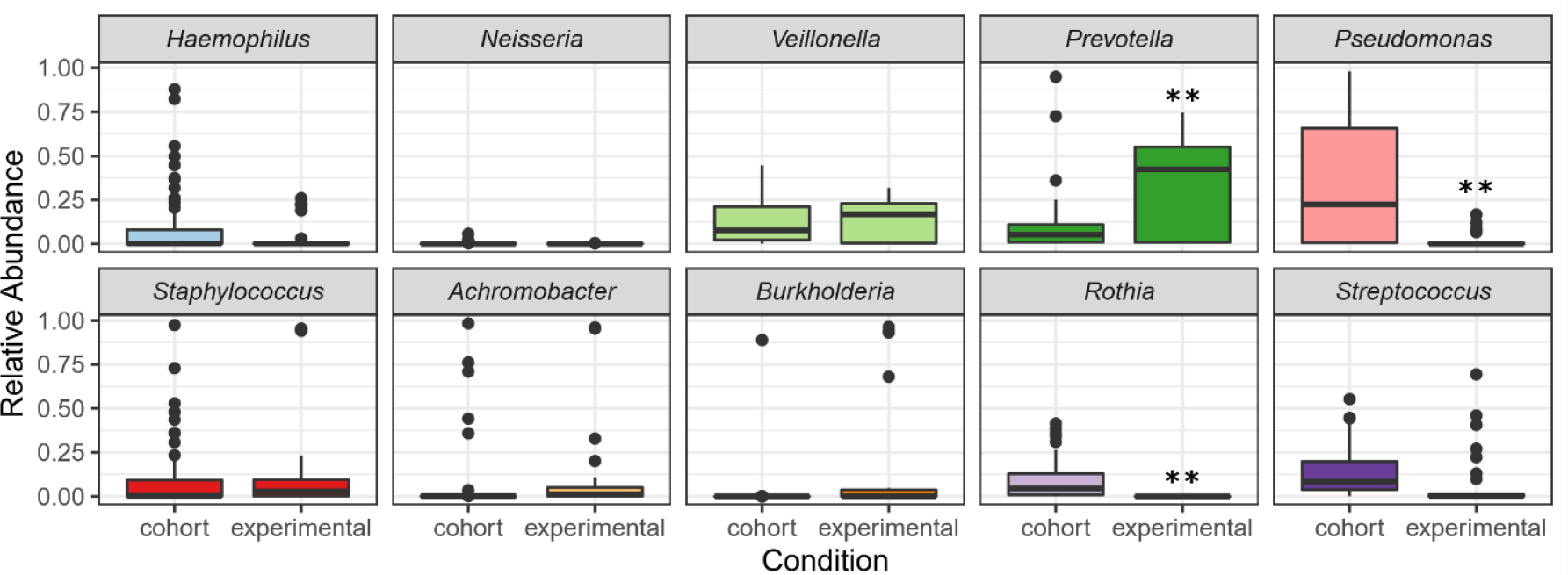
Most experimental taxa fall within the range of clinically observed relative frequencies. The relative abundances of taxa in synthetic microbiome end points (30 samples) compared to 77 clinical cohort observations^57^. The box represents the interquartile range (from 25%-75% of samples) with the horizonal line at the median. Outliers are represented as dots (two-tailed t-test with Bonferroni multiple testing correction: * *p* < 0.05, ** *p* < 0.01).

## Discussion

Our results show that in the absence of antibiotic perturbations, our defined 10-species synthetic CF microbiome community follows a highly repeatable path to a stable community composition (Figures 1, 2, 6). In contrast, antibiotic treatments resulted in diverse trajectories leading to multiple stable end states (Figures 3, 4, 6) that are dispersed throughout a broader space of observed CF community structures including pathogen dominant states (Figure 6).

Our results demonstrate substantial community-scale impacts of antibiotic perturbation, but leave open questions of mechanism. To begin addressing this question, we assessed the extent to which standard antibiotic resistance data (Table S2) are predictive of species presence or absence under antibiotic perturbation in a community context (Table 1). Our results highlight that antibiotics generally enrich for drug resistant taxa and drug-resistant pathogens in particular (Figure 5A, 5B). Yet, MIC data often fails to predict individual species presence (Figure 5C), with predictions showing both false positives (e.g., *B. cenocepacia* under tobramycin, ciprofloxacin or the absence of antibiotics) and false negatives (e.g., *S. aureus* under meropenem). This failure echoes clinical experience, where antibiotic susceptibility testing has been repeatedly shown to not correlate with clinical outcomes of antibiotic treatment^36^.

A simple general explanation for departures from the ecological filter hypothesis is the presence of significant ecological interactions among species. Under this framework, false positives are evidence for suppressive interactions, suggesting that for example *S. aureus* fails to grow in the antibiotic free environment because it is out-competed by one or more of the other taxa. Conversely, false negatives are evidence of facilitation, suggesting for instance that the ability of *S. aureus* to grow in an otherwise lethal dose of meropenem is due to facilitation from another species in the community. Here a likely candidate is the initial presence of *B. cenocepacia*, which encodes multiple β-lactamase enzymes^89^ that are potentially capable of degrading meropenem and therefore may enable *S. aureus* to grow in this environment. Figure 5C indicates an emergence of faciliatory interactions only under antibiotic exposure. This pattern is consistent with the stress-gradient hypothesis, which predicts that positive species interactions are enhanced under environmental stress^90–92^.

While the combination of antibiotic resistance and species interactions is a candidate explanation for our results, other factors are potentially at play. First, we note the important caveat that the MIC estimates were derived using standard growth-promoting rich culture assays, which are known to generate estimates that tend to under-estimate the resistance of cells under more physiologically relevant conditions^93, 94^. If our MICs are under-estimates of resistance, then we would anticipate more ‘false positive’ evidence of competition in our experimental community. A second possibility for divergent results is the presence of physiological or evolutionary adaptation to the community conditions, across the 10 days of serial passaging. The stability and repeatability across replicates in Figure 1 argue against a major role for genetic evolution in steering community dynamics – consistent with recent work on the suppressive impact of community interactions on bacterial evolution^95^.

In order to develop an experimentally tractable model, we made a number of choices regarding specific experimental conditions (nutrients, initial community structure) that likely influenced our specific results. The healthy lung is evidently an oxygen rich environment, however during the course of tissue degradation in the CF airways, the sputum environment can become oxygen deprived due to the combined forces of mucus plugs, along with oxygen consumption by immune cells and microbes^58, 59, 96^. To capture an oxygen stressed environment, we performed our experiments under static anaerobic conditions that were only subjected to oxygenation during bench passaging every 48 h. While all bacteria in the community are capable of either fermentation, anaerobic respiration, or both, the largely anaerobic condition represents a potential to bias the results towards strictly anaerobic bacteria. Our clinical benchmarking exercise indicates that the distribution of oxygen exposures in the clinic is less biased towards anaerobic conditions, as our three taxon ‘misses’ (Figure 7) consisted of over-representation of an anaerobe (*P. melaninogenica*) and under-representation of two aerobes (*P. aeruginosa*, *R. mucilaginosa*). This pattern is also consistent with recent transcriptomic analyses of *P. aeruginosa* from CF sputum, highlighting a transcriptional response indicative of reduced oxygen, but not necessarily anaerobic conditions^97^. In future work we will investigate synthetic community dynamics in static communities with partial exposure to room air, following recent experimental *ex vivo* (patient sputum) models^98, 99^.

Turning to our choices regarding synthetic community composition, by focusing on the most abundant bacterial taxa, we ignored the potential for rare keystone species to shape community dynamics^100^. We also overlooked the potential importance of interactions among strains within each species^101, 102^. Concerning specific strain choices, Figure 2 illustrates that replacing *P. aeruginosa* PDO300 with an otherwise isogenic non-mucoid strain (PAO1) produces little dynamical change. However other studies in different environmental contexts have demonstrated substantial dependency of interactions on strain identity^103, 104^, leaving open the importance of specific strain identities in governing community outcomes. More broadly, we did not include other potentially critical players in the lung microbiome, spanning human epithelial and immune cells, fungal species, and viruses of all the above. We note that our experimental platform is amenable to the addition of these players in future controlled experiments.

Finally, our results leave open the importance of initial community composition. Our initial community state was positive for all 10 species, including all major pathogens, and is representative of the average metacommunity^105^ of a population of people with CF^57^. While each of these species is common in people with CF, they are not all commonly found together – in particular, the pathogens tend to be more segregated across patients^54, 57^. Despite our starting-point being un-representative of individual patient profiles (see Figure 6 for visualization of distance from inoculum to clinical community structures), our experiments show movement towards a diverse set of clinically observed states with greater dominance by fewer pathogens (Figures 3, 4, 6). Assessing the importance of initial conditions in determining final outcome is a worthwhile avenue for future study, as other studies have demonstrated that differing initial proportions of the same set of strains can lead to differing dynamical outcomes, particularly in cases of interference competition^106^.

Our results demonstrate the power of a model 10-species system for the study of chronic lung infection dynamics, validated against data from people with CF (Figure 6). This model provides a platform to assess the community ecological impacts of currently deployed antibiotic treatments (Figures 2-4, 6) and novel treatments – from different compounds to different strategies of their implementation. Current practice is to ‘hit hard’ with an antibiotic that is effective against a target pathogen^107^. In the context of our model community, detecting drug susceptible *P. aeruginosa* would typically trigger combination treatments that lead in our example to rapid emergence of more dominant and more resistant pathogen replacements (Figures 3-5)^108^. One way to improve on this picture is to run community-scale resistance diagnostics, and in turn use this diagnostic information to optimize antibiotic (and probiotic) choices^109^. While simple in outline, identifying optimal treatment choices in the context of complex multi-species communities poses a substantial computational and experimental challenge.

## Materials and Methods

### Bacterial strains

Table 1 outlines the specific strains in our 10-species community. Species choices were initially informed based on our previous study of a 77-person CF cohort^57^. Our 10 species represent the most abundant genera from our 16S rDNA analyses (together accounting for over 85% of reads). Note that these species are collectively representative of the ‘metacommunity’ (the community of communites^105^) of microbes across a population of people with CF, and are not necessarily representative of individual community states. We view this metacommunity as the menu of organisms from which individual communities are sampled.

To guide our experimental species choices, we turned to existing CF metagenome sequencing data^110^, which provided high confidence for all but one of our species calls (Table 1). The exception is *Streptococcus*, where reads are distributed across a range of species. We chose *S. mitis* because it is present in sputum metagenomic profiles^110^, and it is an experimentally tractable organism that is typically considered to be non-pathogenic^111^. Within each species, we focused on well-characterized reference strains, as far as these were available, including American Type Culture Collection (ATCC) strains. For the dominant pathogen *P. aeruginosa* (PA), we used both the reference strain PAO1 and its mucoid derivative PDO300^112^. Our default experimental choice is PDO300, as this strain better reflects the mucoid phenotype prevalent in chronic CF^112, 113^.

### Community growth medium

Our Artificial Sputum Medium (ASM) is based on the benchmarked synthetic CF sputum medium 2 (SCFM2^55, 56^), but with differences in the preparation of the mucin and DNA macro-molecules. Specifically, mucins were ethanol washed and autoclaved (not UV sterilized, due to larger volume requirements), and the entire medium was filter sterilized following addition of DNA. Given the potential for differences in preparation methods to impact the results, we refer to our medium under the more generic name of ASM to underline these differences from the reference recipe for SCFM2^55, 56^.

### Bacterial pre-culture and community construction

Before the experiment, all bacterial strains were revived from frozen stocks by streaking on rich media agar plates (chocolate or BHI agar, depending on the species, see Table S1) and cultured at 37°C for 48 hours (microaerophilically (for *H. influenzae* and *N. subflava*) or anaerobically (for *P. melaninogenica* and *V. parvula*, in GasPak jars). Five colonies were then picked from each plate and used to inoculate specific monoculture rich medium, which was cultured for a further 48 h; specific culture conditions are detailed in Table S1.

The bacterial cultures were then washed in a defined ASM buffer base (ASM minus all carbon sources), OD_600_ values were measured with a Hidex plate reader (Hidex Oy, Finland) and adjusted to 0.5 for each species and diluted 10-fold in ASM. These standardized bacterial dilutions of equal volume were mixed and antibiotic stocks were added according to the experimental design for each treatment. The bacterial mixtures (plus antibiotics, dependent on treatment) were homogenized with pipetting, then divided into five replicates of 2 ml each in 24-well plates. An additional 0.5 ml of the initial inoculum mixture was stored at - 80°C to assess community composition at time zero by subsequent genomic analysis.

### Treatments and passaging

To measure the impact of exposure to antibiotics, we tested three antibiotics that are widely used in CF therapy: tobramycin (5 μg / ml); meropenem (15 μg / ml); ciprofloxacin (2.5 μg / ml), and two widely used combinations; tobramycin and meropenem; tobramycin and ciprofloxacin (adding the concentrations above). The specific concentrations used reflect measurements of antibiotic concentrations in CF sputum^65, 114, 115^. All experiments were performed with 5 replicates of 2 mL cultures in 24-well plates cultured at 37°C in anaerobic GasPak jars. Every 48 h, bacterial cultures were mixed by pipetting, and 10% of the volume was transferred to fresh ASM (with fresh antibiotics as defined by the treatment). 0.5 ml of the culture was stored at each passage at -80°C for later DNA purification and amplicon sequencing. Each experimental line was maintained for 5 passages (10 days).

To assess the role of pathogen identity in community composition, the *P. aeruginosa* and *S. aureus* composition of the standard community was altered. The 5 resulting communities were either free of both pathogens or contained either *P. aeruginosa* PAO1 or mucoid *P. aeruginosa* PDO300^112^, in either case with or without *S. aureus*. These experiments were done in the absence of antibiotics, but otherwise with the same conditions as above.

### 16S rDNA sequencing and qPCR

DNA purification, sequencing, and qPCR were performed by MR DNA Lab (Shallowater, TX). Briefly: DNA was purified from sputum homogenate after mechanical lysis with the MoBio Power Soil kit (MoBio, Carlsbad, CA). The 16S V4 region of the resulting DNA was amplified with 515F and 806R primers incorporating the barcode in the forward primer and subjected to Illumina sequencing^116^. The sequence data were generated in a total of 6 MiSeq runs. Total 16S abundance in each sample was determined by qPCR using standard 515F/806R primers^116^.

### 16S rDNA sequence analysis

To generate taxa counts from the sequence data, we processed each run independently and combined the results. Across the 6 sequencing runs, a total of 15,347,658 sequence reads were generated, with a median of 59,686 sequences per sample (minimum 22,707, maximum 126,680). All sequence processing was done through QIIME2 2019.10.0. Unless otherwise noted, we left parameters as defaults based on the Moving Pictures workflow. Samples were demultiplexed using the cutadapt plugin in QIIME2. We found that some of the barcode sequences were also found in the 16S region of several taxa. To mitigate this confounder, we removed from each metadata file the first four nucleotides in the 515F primer and added it to the barcode. For example, the barcode “GAGATGTG” was remapped as “GAGATGTGGTGC” and the primer became “CAGCMG…”.

### 16S rDNA sequencing and qPCR

Reads were denoised using the deblur plugin, and resulting sequences were trimmed to 250 bp. Taxonomic assignments were classified against the greengenes 16S database. Some assignments were not possible at a level of genus resolution, so we interpreted reads mapping to “o Lactobacillales” to “g Streptococcus”, “f Burkholderiaceae” as “g Burkholderia”, and “f Pseudomonadaceae” as “g_Pseudomonas”. Finally, for each sample we removed spurious (and rare) taxon calls that did not map onto our experimentally defined communities. Sequence data have been deposited to the SRA (Accession # deposit pending). The analysis pipeline is available on GitHub (GitHub pending).

Absolute abundances were determined by proportion of the total 16S count and then normalized to species-specific 16S rDNA copy counts^57, 117^.

### Statistical analyses

All analyses and plots used the R programming language^118, 119^. Tables and scripts can be found at (GitHub pending). A nonparametric Wilcoxon rank sum test was used to test for differences in absolute species abundances across experimental conditions, using a two-tailed test for change in abundance and a one-tailed test to assess competitive release (testing for increases only). A t-test with a Bonferroni multiple testing correction was performed to compare relative abundances of the 10 species under experimental conditions with clinical samples from the 77-patient cohort. To compare experimental treatments (and clinical benchmark data) at a community scale, we calculated ANOSIM R values on Bray-Curtis dissimilarity matrices for each treatment using the vegan package^74–76^. The R statistic is a ratio of within-treatment differences to between-treatment differences on a scale of -1 to 1, where a value of 1 would mean that all dissimilarity is between treatments, indicating completely different communities.

To visualize community-scale differences we constructed ordination plots for combined clinical and experimental compositional data. Clinical and experimental observations were center-log-transformed first^120, 121^, then standardized before principal component analysis^118, 121^.

## Acknowledgments

We thank the Brown lab, the Center for Microbial Dynamics and Infection, and Drs. Sylvie Estrela, Ellinor Alseth, and John LiPuma for valuable discussion and feedback. We thank Drs. Joanna Goldberg and Bob Jerris for providing bacteria strains. This work was supported in part by CDC contracts BAA 2016-N-17812, BAA 2017-OADS-01, and 75D30120C-09782 to SPB, and CFF awards BROWN19I0 and BROWN21P0 to SPB, GURNEY20F0 to JRG and DAVIS21H0 to JDD.

## Supplemental Figures

**Fig. S1.**
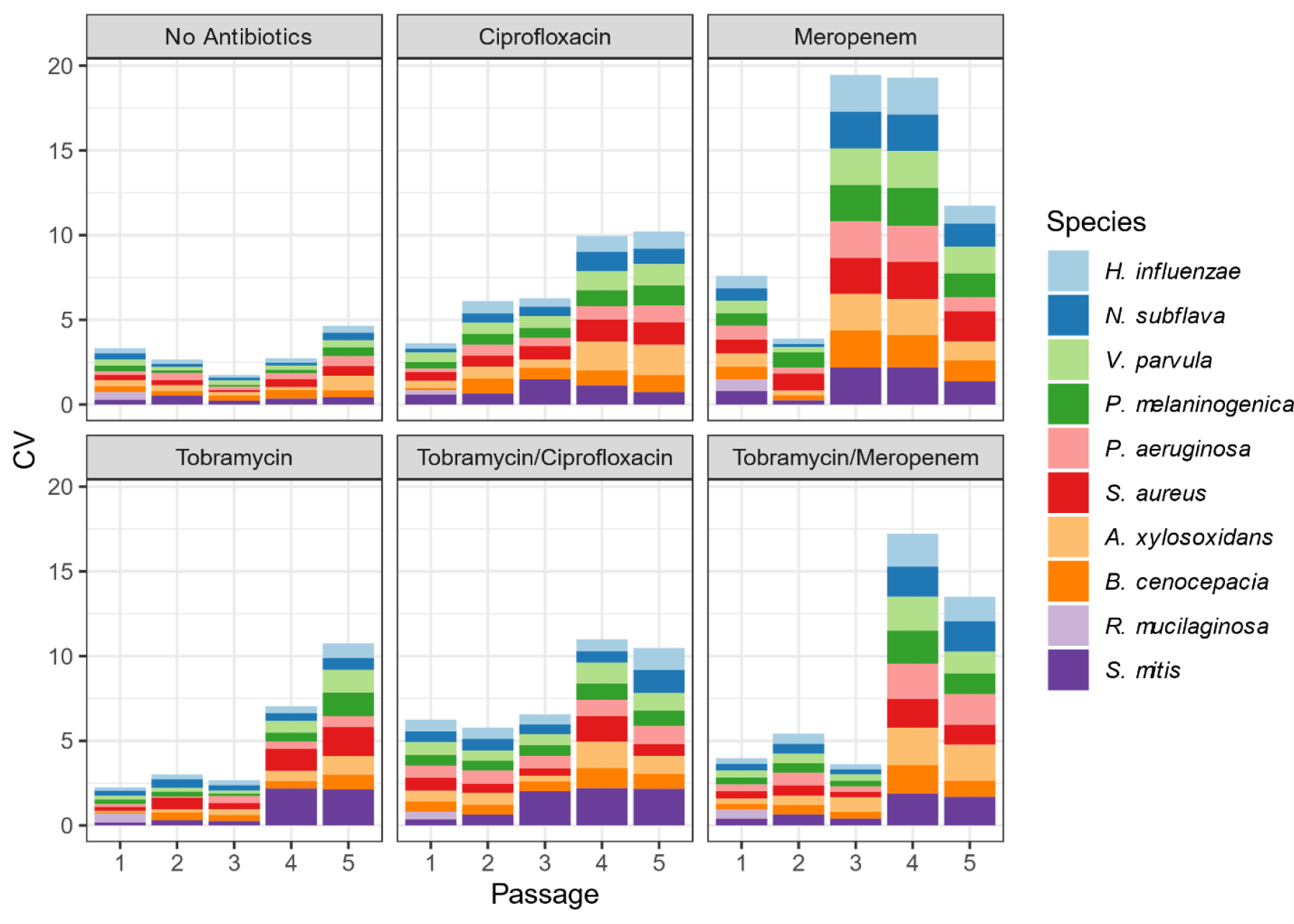
Coefficient of Variance (CV) in species abundances, across replicates. CVs under different treatments through time. Stacked bars represent the CV of each individual species. CV is calculated as the standard deviation (across replicates) at each passage divided by its mean.

**Fig. S2.**
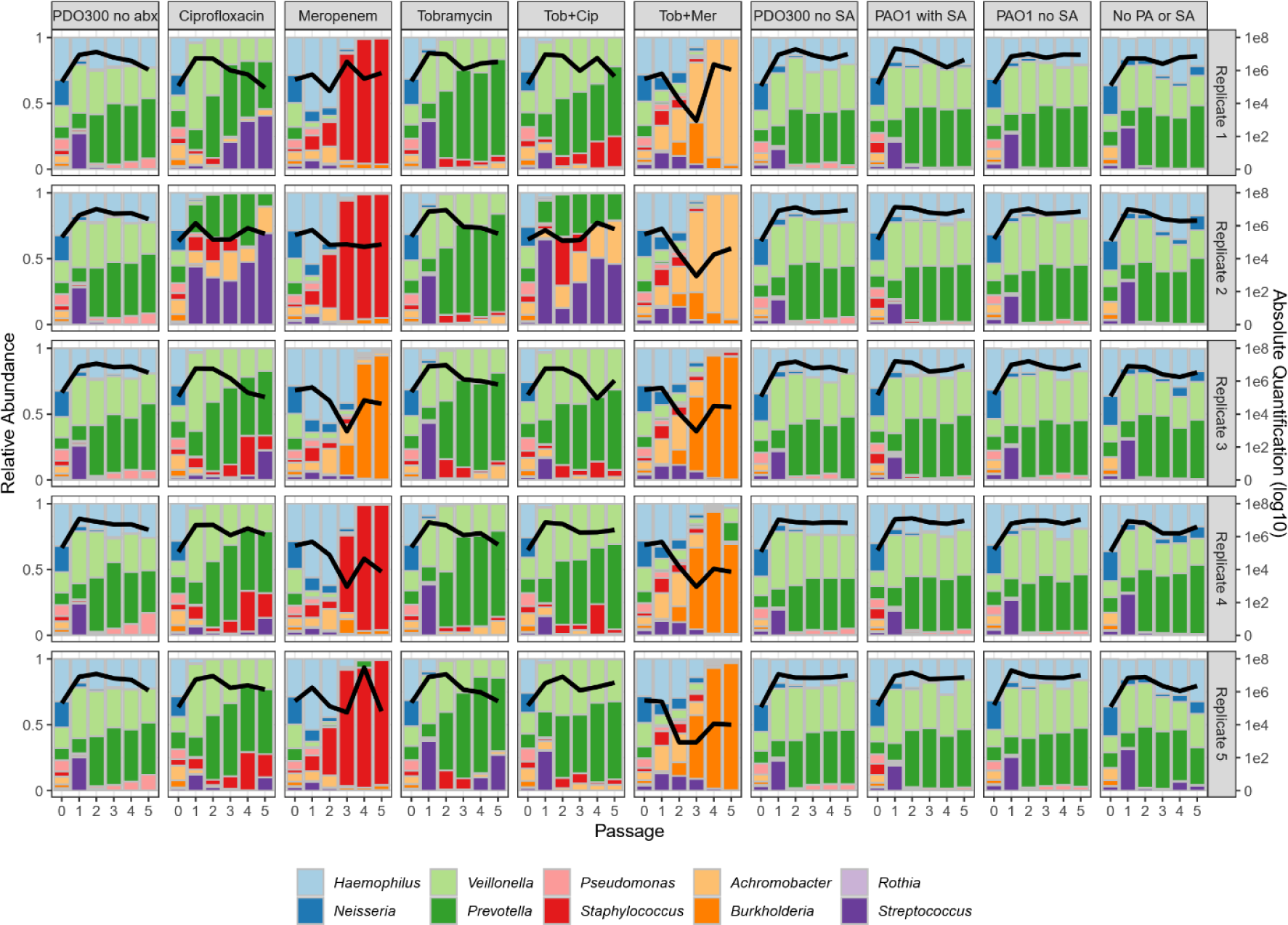
Compositional and total abundances across all treatments (columns) and replicates (rows). For details, see legend of Figure 1.

**Fig. S3.**
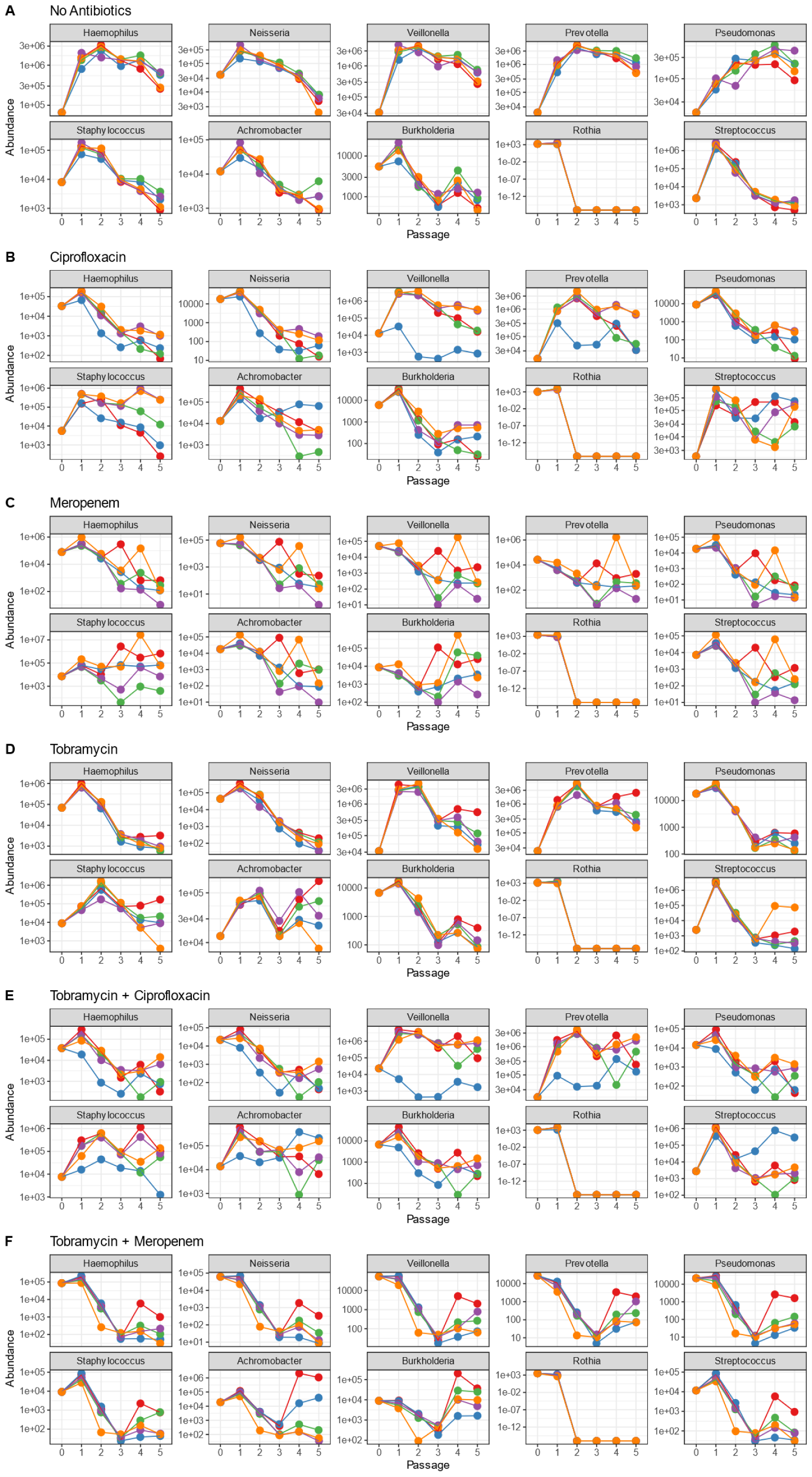
Temporal absolute abundances across all treatments (panels A-F) and all taxa (sub-panels). Data are plotted for each species under each antibiotic condition (A-F). The X-axis represents passage and Y-axis represents absolute abundance per ml.

**Fig. S4.**
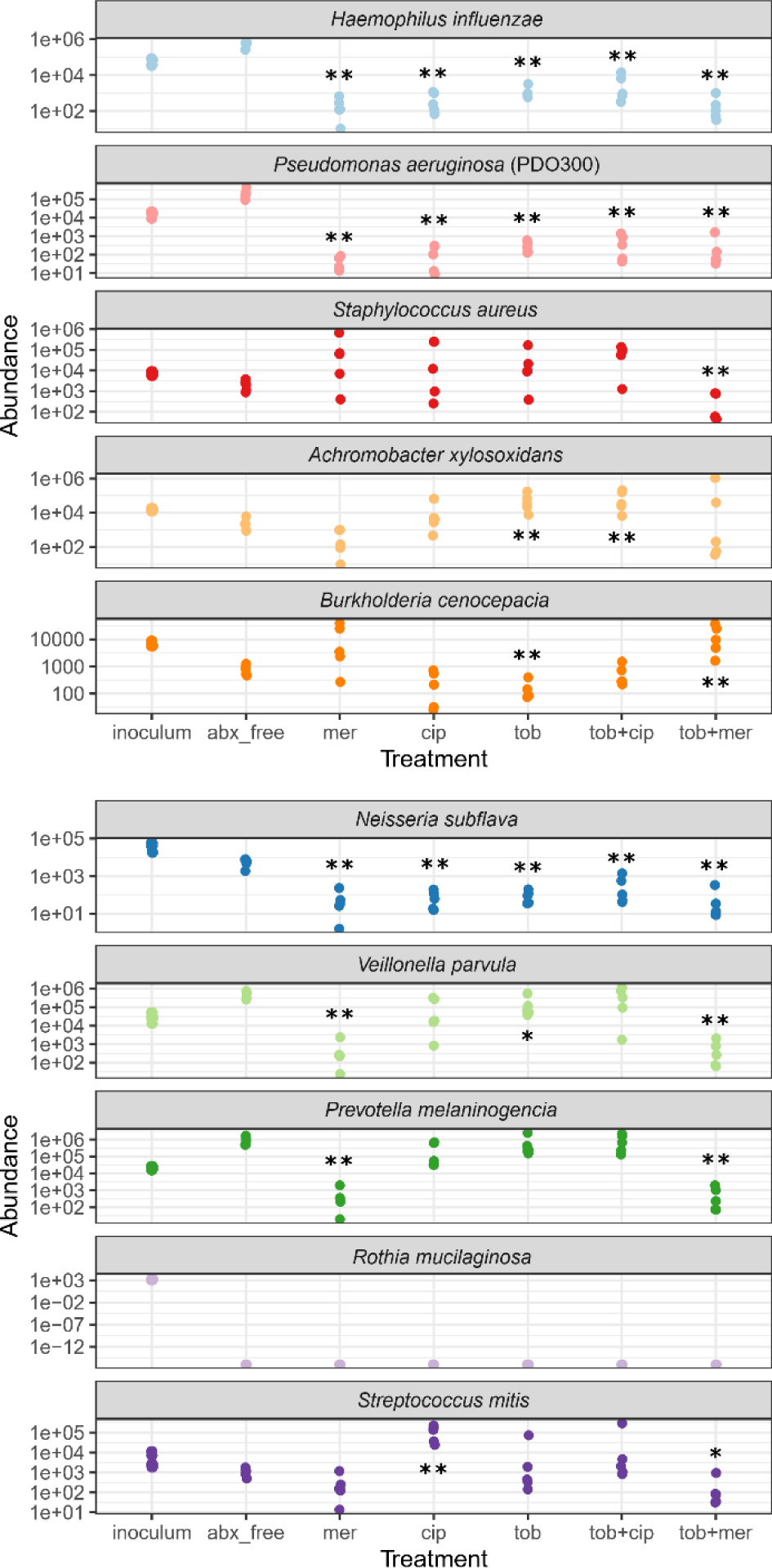
Absolute microbe densities across antibiotic exposures. Each dot corresponds to an individual replicate of species-specific initial (inoculum) and final time-point absolute density under defined antibiotic treatments (data redrawn from Figure 3). abx_free = no antibiotic condition, mer = meropenem, cip = ciprofloxacin, tob = tobramycin. Asterisks denote significantly higher/lower final densities in presence of antibiotic, compared to antibiotic-free controls (two-tailed Wilcoxon test, * *p* < 0.05, ** *p* < 0.01).

**Fig. S5.**
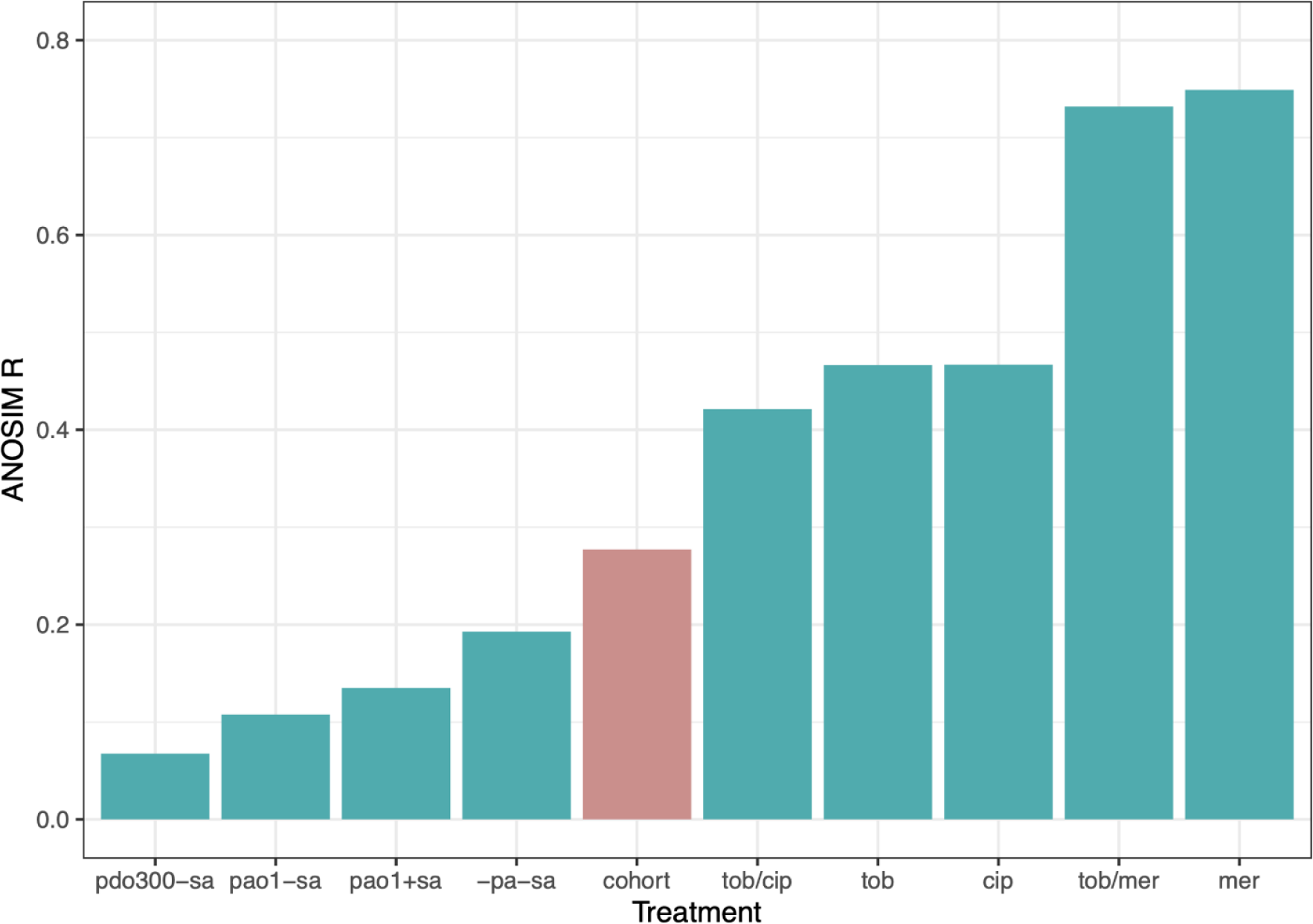
Differences in community structures across experimental treatments and clinical data. The ANOSIM R statistic captures the ratio of between-group to within-group variances; as R approaches 1, more variance is found between groups than within groups^75^. Green boxes are comparisons between absolute abundances of the ‘no drug’ reference treatment (PDO300 with SA, Figure 1) and experimental manipulations (pathogen treatments, Figure 2, and drug treatments, Figure 3). The pink box is the comparison between relative abundances of all experimental conditions (Figures 1, 2, 3) and all clinical conditions. Passage 0 (inoculum) is omitted from the analyses. All R values are significant with *p* < 0.05 (permutation test).

**Table S1.** Monoculture pre-culture conditions. The atmospheric environment specifies the oxygenation used for both the agar plate (Brain Heart Infusion (BHI) or chocolate agar) and liquid culture steps. The liquid medium supplements had the following concentrations: hemin, 15 mg / L; NAD, 15 mg / L; vitamin K1, 1 mg / L; L-lactate, 50 mM. Note that in subsequent experiments we simplified the protocol so that all bacteria were cultured first on chocolate agar plates, and then in a common medium of TSYE supplemented with hemin, NAD, vitamin K, and lactic acid

| Genus | Species | Oxygen tolerance | Liquid medium | Agar plates |
| --- | --- | --- | --- | --- |
| *Pseudomonas* | *aeruginosa* | Aerobic | TSYE | BHI |
| *Staphylococcus* | *aureus* | Aerobic | TSYE | BHI |
| *Achromobacter* | *xylosoxidans* | Aerobic | TSYE | BHI |
| *Haemophilus* | *influenzae* | Microaerophilic | TSYE+hemin+NAD | Chocolate agar |
| *Streptococcus* | *Mitis* | Aerobic | TSYE | BHI |
| *Rothia* | *mucilaginosa* | Aerobic | TSYE | Chocolate agar |
| *Burkholderia* | *cenocepacia* | Aerobic | TSYE | BHI |
| *Neisseria* | *subflava* | Microaerophilic | TSYE | Chocolate agar |
| *Prevotella* | *melaninogenica* | Anaerobic | TSYE+hemin+VK1 | Chocolate agar |
| *Veillonella* | *parvula* | Anaerobic | TSYE+lactate | Chocolate agar |

**Table S2.**
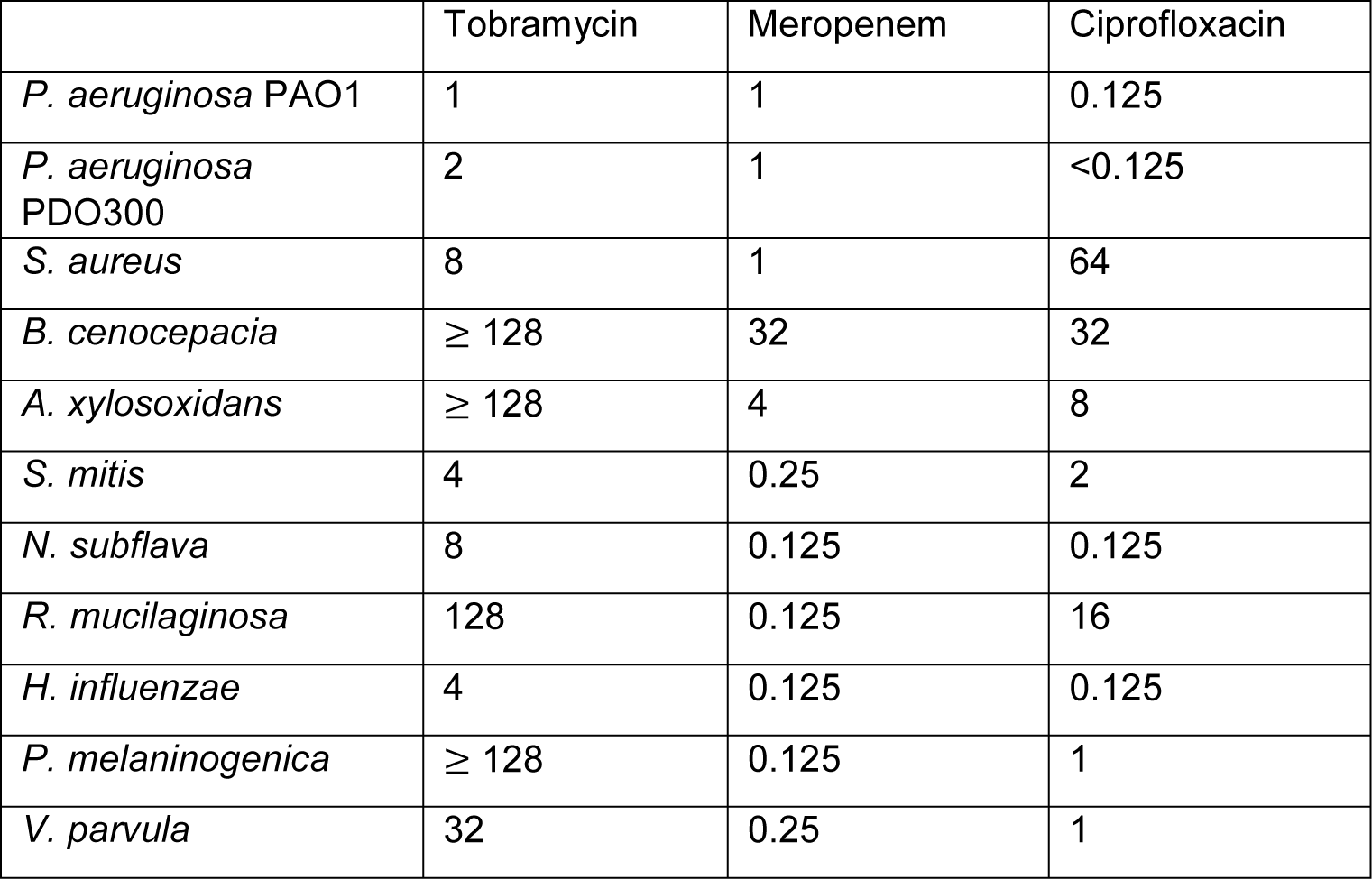
Antibiotic susceptibility in rich medium. Minimal Inhibitory Concentrations (MICs, in μg / ml) of synthetic community members were determined in rich medium.

## References

1. Laxminarayan, R. et al. Antibiotic resistance - the need for global solutions. The Lancet Infectious Diseases 13, 1057–1098, doi:10.1016/S1473-3099(13)70318-9 (2013).

2. Filkins, L. M. & O’Toole, G. A. Cystic Fibrosis Lung Infections: Polymicrobial, Complex, and Hard to Treat. PLoS pathogens 11, e1005258, doi:10.1371/journal.ppat.1005258 (2015).

3. Siddiqui, A. R. & Bernstein, J. M. Chronic wound infection: facts and controversies. Clinics in dermatology 28, 519–526, doi:10.1016/j.clindermatol.2010.03.009 (2010).

4. Young, D., Hussell, T. & Dougan, G. Chronic bacterial infections: living with unwanted guests. Nature immunology 3, 1026–1032, doi:10.1038/ni1102-1026 (2002).

5. Guest, J. F. et al. Health economic burden that different wound types impose on the UK’s National Health Service. International wound journal 14, 322–330, doi:10.1111/iwj.12603 (2017).

6. Bjarnsholt, T. et al. The in vivo biofilm. Trends Microbiol 21, 466–474, doi:10.1016/j.tim.2013.06.002 (2013).

7. Darch, S. E. et al. Phage Inhibit Pathogen Dissemination by Targeting Bacterial Migrants in a Chronic Infection Model. mBio 8, doi:10.1128/mBio.00240-17 (2017).

8. Kragh, K. N. et al. Polymorphonuclear leukocytes restrict growth of *Pseudomonas aeruginosa* in the lungs of cystic fibrosis patients. Infect Immun 82, 4477–4486, doi:10.1128/iai.01969-14 (2014).

9. Stacy, A., McNally, L., Darch, S. E., Brown, S. P. & Whiteley, M. The biogeography of polymicrobial infection. Nat Rev Microbiol 14, 93–105, doi:10.1038/nrmicro.2015.8 (2016).

10. Henke, M. O. & Ratjen, F. Mucolytics in cystic fibrosis. Paediatric respiratory reviews 8, 24–29, doi:10.1016/j.prrv.2007.02.009 (2007).

11. Perez-Vilar, J. & Boucher, R. C. Reevaluating gel-forming mucins’ roles in cystic fibrosis lung disease. Free radical biology & medicine 37, 1564–1577, doi:10.1016/j.freeradbiomed.2004.07.027 (2004).

12. Surette, M. G. The cystic fibrosis lung microbiome. Annals of the American Thoracic Society 11 **Suppl 1**, S61–65, doi:10.1513/AnnalsATS.201306-159MG (2014).

13. Yonker, L. M., Cigana, C., Hurley, B. P. & Bragonzi, A. Host-pathogen interplay in the respiratory environment of cystic fibrosis. Journal of Cystic Fibrosis 14, 431–439, doi:doi.org/10.1016/j.jcf.2015.02.008 (2015).

14. CFF. Cystic Fibrosis Foundation patient registry 2019 annual data report., (Cystic Fibrosis Foundation, 2019).

15. Fodor, A. A. et al. The adult cystic fibrosis airway microbiota is stable over time and infection type, and highly resilient to antibiotic treatment of exacerbations. PloS one 7, e45001, doi:10.1371/journal.pone.0045001 (2012).

16. Frayman, K. B., Armstrong, D. S., Grimwood, K. & Ranganathan, S. C. The airway microbiota in early cystic fibrosis lung disease. Pediatric pulmonology 52, 1384–1404, doi:10.1002/ppul.23782 (2017).

17. Huang, Y. J. & LiPuma, J. J. The Microbiome in Cystic Fibrosis. Clinics in chest medicine 37, 59–67, doi:10.1016/j.ccm.2015.10.003 (2016).

18. Lucas, S. K., Yang, R., Dunitz, J. M., Boyer, H. C. & Hunter, R. C. 16S rRNA gene sequencing reveals site-specific signatures of the upper and lower airways of cystic fibrosis patients. Journal of Cystic Fibrosis 17, 204–212, doi:doi.org/10.1016/j.jcf.2017.08.007 (2018).

19. Filkins, L. M. et al. Prevalence of *streptococci* and increased polymicrobial diversity associated with cystic fibrosis patient stability. Journal of bacteriology 194, 4709–4717, doi:10.1128/jb.00566-12 (2012).

20. Caverly, L. J. & LiPuma, J. J. Good cop, bad cop: anaerobes in cystic fibrosis airways. The European respiratory journal 52, doi:10.1183/13993003.01146-2018 (2018).

21. Acosta, N. et al. Sputum microbiota is predictive of long-term clinical outcomes in young adults with cystic fibrosis. Thorax 73, 1016–1025, doi:10.1136/thoraxjnl-2018-211510 (2018).

22. Coburn, B. et al. Lung microbiota across age and disease stage in cystic fibrosis. Scientific reports 5, 10241, doi:10.1038/srep10241 (2015).

23. Muhlebach, M. S. et al. Initial acquisition and succession of the cystic fibrosis lung microbiome is associated with disease progression in infants and preschool children. PLoS pathogens 14, e1006798, doi:10.1371/journal.ppat.1006798 (2018).

24. Zhao, J. et al. Decade-long bacterial community dynamics in cystic fibrosis airways. Proceedings of the National Academy of Sciences of the United States of America 109, 5809–5814, doi:10.1073/pnas.1120577109 (2012).

25. Adamowicz, E. M., Flynn, J., Hunter, R. C. & Harcombe, W. R. Cross-feeding modulates antibiotic tolerance in bacterial communities. The ISME journal 12, 2723–2735, doi:10.1038/s41396-018-0212-z (2018).

26. Flynn, J. M., Niccum, D., Dunitz, J. M. & Hunter, R. C. Evidence and Role for Bacterial Mucin Degradation in Cystic Fibrosis Airway Disease. PLoS pathogens 12, e1005846, doi:10.1371/journal.ppat.1005846 (2016).

27. Goddard, A. F. et al. Direct sampling of cystic fibrosis lungs indicates that DNA-based analyses of upper-airway specimens can misrepresent lung microbiota. Proceedings of the National Academy of Sciences of the United States of America 109, 13769–13774, doi:10.1073/pnas.1107435109 (2012).

28. Jorth, P. et al. Direct Lung Sampling Indicates That Established Pathogens Dominate Early Infections in Children with Cystic Fibrosis. Cell reports 27, 1190–1204.e1193, doi:10.1016/j.celrep.2019.03.086 (2019).

29. Rogers, G. B. et al. Use of 16S rRNA gene profiling by terminal restriction fragment length polymorphism analysis to compare bacterial communities in sputum and mouthwash samples from patients with cystic fibrosis. Journal of clinical microbiology 44, 2601–2604, doi:10.1128/jcm.02282-05 (2006).

30. Hogan, D. A. et al. Analysis of Lung Microbiota in Bronchoalveolar Lavage, Protected Brush and Sputum Samples from Subjects with Mild-To-Moderate Cystic Fibrosis Lung Disease. PloS one 11, e0149998, doi:10.1371/journal.pone.0149998 (2016).

31. Muhlebach, M. S. et al. Anaerobic bacteria cultured from cystic fibrosis airways correlate to milder disease: a multisite study. The European respiratory journal 52, doi:10.1183/13993003.00242-2018 (2018).

32. Zemanick, E. T. et al. Assessment of airway microbiota and inflammation in cystic fibrosis using multiple sampling methods. Annals of the American Thoracic Society 12, 221–229, doi:10.1513/AnnalsATS.201407-310OC (2015).

33. Lu, J. et al. Parallel Analysis of Cystic Fibrosis Sputum and Saliva Reveals Overlapping Communities and an Opportunity for Sample Decontamination. mSystems 5, doi:10.1128/mSystems.00296-20 (2020).

34. Chmiel, J. F. et al. Antibiotic management of lung infections in cystic fibrosis. I. The microbiome, methicillin-resistant Staphylococcus aureus, gram-negative bacteria, and multiple infections. Annals of the American Thoracic Society 11, 1120–1129, doi:10.1513/AnnalsATS.201402-050AS (2014).

35. Waters, V. Chronic Antibiotic Use in Cystic Fibrosis: A Fine Balance. Annals of the American Thoracic Society 15, 667–668, doi:10.1513/AnnalsATS.201803-172ED (2018).

36. Somayaji, R. et al. Antimicrobial susceptibility testing (AST) and associated clinical outcomes in individuals with cystic fibrosis: A systematic review. Journal of cystic fibrosis : official journal of the European Cystic Fibrosis Society 18, 236–243, doi:10.1016/j.jcf.2019.01.008 (2019).

37. Kidd, T. J. et al. Defining antimicrobial resistance in cystic fibrosis. Journal of cystic fibrosis : official journal of the European Cystic Fibrosis Society 17, 696–704, doi:10.1016/j.jcf.2018.08.014 (2018).

38. Darch, S. E. et al. Recombination is a key driver of genomic and phenotypic diversity in a *Pseudomonas aeruginosa* population during cystic fibrosis infection. Scientific reports 5, 7649, doi:10.1038/srep07649 (2015).

39. Jorth, P. et al. Regional Isolation Drives Bacterial Diversification within Cystic Fibrosis Lungs. Cell host & microbe 18, 307–319, doi:10.1016/j.chom.2015.07.006 (2015).

40. Waters, V. J. et al. Reconciling Antimicrobial Susceptibility Testing and Clinical Response in Antimicrobial Treatment of Chronic Cystic Fibrosis Lung Infections. Clinical infectious diseases : an official publication of the Infectious Diseases Society of America 69, 1812–1816, doi:10.1093/cid/ciz364 (2019).

41. Cuthbertson, L. et al. Respiratory microbiota resistance and resilience to pulmonary exacerbation and subsequent antimicrobial intervention. The ISME journal 10, 1081–1091, doi:10.1038/ismej.2015.198 (2016).

42. Price, K. E. et al. Unique microbial communities persist in individual cystic fibrosis patients throughout a clinical exacerbation. Microbiome 1, 27, doi:10.1186/2049-2618-1-27 (2013).

43. Hahn, A. et al. Changes in microbiome diversity following beta-lactam antibiotic treatment are associated with therapeutic versus subtherapeutic antibiotic exposure in cystic fibrosis. Scientific reports 9, 2534, doi:10.1038/s41598-019-38984-y (2019).

44. Smith, D. J. et al. Pyrosequencing reveals transient cystic fibrosis lung microbiome changes with intravenous antibiotics. The European respiratory journal 44, 922–930, doi:10.1183/09031936.00203013 (2014).

45. Zemanick, E. T. et al. Inflammation and airway microbiota during cystic fibrosis pulmonary exacerbations. PloS one 8, e62917, doi:10.1371/journal.pone.0062917 (2013).

46. Nelson, M. T. et al. Maintenance tobramycin primarily affects untargeted bacteria in the CF sputum microbiome. Thorax 75, 780–790, doi:10.1136/thoraxjnl-2019-214187 (2020).

47. Taccetti, G. et al. Early antibiotic treatment for *Pseudomonas aeruginosa* eradication in patients with cystic fibrosis: a randomised multicentre study comparing two different protocols. Thorax 67, 853–859, doi:10.1136/thoraxjnl-2011-200832 (2012).

48. Morley, V. J., Woods, R. J. & Read, A. F. Bystander Selection for Antimicrobial Resistance: Implications for Patient Health. Trends Microbiol 27, 864–877, doi:10.1016/j.tim.2019.06.004 (2019).

49. Hotterbeekx, A., Kumar-Singh, S., Goossens, H. & Malhotra-Kumar, S. In vivo and In vitro Interactions between *Pseudomonas aeruginosa* and *Staphylococcus* spp. Frontiers in cellular and infection microbiology 7, 106, doi:10.3389/fcimb.2017.00106 (2017).

50. Limoli, D. H. et al. *Pseudomonas aeruginosa* Alginate Overproduction Promotes Coexistence with *Staphylococcus aureus* in a Model of Cystic Fibrosis Respiratory Infection. mBio 8, doi:10.1128/mBio.00186-17 (2017).

51. Nguyen, A. T. & Oglesby-Sherrouse, A. G. Interactions between *Pseudomonas aeruginosa* and *Staphylococcus aureus* during co-cultivations and polymicrobial infections. Applied microbiology and biotechnology 100, 6141–6148, doi:10.1007/s00253-016-7596-3 (2016).

52. Vandeplassche, E., Coenye, T. & Crabbé, A. Developing selective media for quantification of multispecies biofilms following antibiotic treatment. PloS one 12, e0187540, doi:10.1371/journal.pone.0187540 (2017).

53. Vandeplassche, E. et al. Antibiotic susceptibility of cystic fibrosis lung microbiome members in a multispecies biofilm. Biofilm 2, 100031, doi:10.1016/j.bioflm.2020.100031 (2020).

54. Jean-Pierre, F., Vyas, A., Hampton, T. H., Henson, M. A. & O’Toole, G. A. One versus Many: Polymicrobial Communities and the Cystic Fibrosis Airway. mBio 12, doi:10.1128/mBio.00006-21 (2021).

55. Palmer, K. L., Mashburn, L. M., Singh, P. K. & Whiteley, M. Cystic fibrosis sputum supports growth and cues key aspects of *Pseudomonas aeruginosa* physiology. Journal of bacteriology 187, 5267–5277, doi:10.1128/JB.187.15.5267-5277.2005 (2005).

56. Turner, K. H., Wessel, A. K., Palmer, G. C., Murray, J. L. & Whiteley, M. Essential genome of *Pseudomonas aeruginosa* in cystic fibrosis sputum. Proceedings of the National Academy of Sciences of the United States of America 112, 4110–4115, doi:10.1073/pnas.1419677112 (2015).

57. Zhao, C. Y. et al. Microbiome data enhances predictive models of lung function in people with cystic fibrosis. The Journal of infectious diseases, doi:10.1093/infdis/jiaa655 (2020).

58. Cowley, E. S., Kopf, S. H., LaRiviere, A., Ziebis, W. & Newman, D. K. Pediatric Cystic Fibrosis Sputum Can Be Chemically Dynamic, Anoxic, and Extremely Reduced Due to Hydrogen Sulfide Formation. mBio 6, e00767, doi:10.1128/mBio.00767-15 (2015).

59. Kolpen, M. et al. Polymorphonuclear leucocytes consume oxygen in sputum from chronic *Pseudomonas aeruginosa* pneumonia in cystic fibrosis. Thorax 65, 57–62, doi:10.1136/thx.2009.114512 (2010).

60. Worlitzsch, D. et al. Effects of reduced mucus oxygen concentration in airway *Pseudomonas* infections of cystic fibrosis patients. The Journal of clinical investigation 109, 317–325, doi:10.1172/jci13870 (2002).

61. Tunney, M. M. et al. Detection of anaerobic bacteria in high numbers in sputum from patients with cystic fibrosis. Am J Respir Crit Care Med 177, 995–1001, doi:10.1164/rccm.200708-1151OC (2008).

62. Wong, K., Roberts, M. C., Owens, L., Fife, M. & Smith, A. L. Selective media for the quantitation of bacteria in cystic fibrosis sputum. J Med Microbiol 17, 113–119, doi:10.1099/00222615-17-2-113 (1984).

63. O’Neill, K. et al. Reduced bacterial colony count of anaerobic bacteria is associated with a worsening in lung clearance index and inflammation in cystic fibrosis. PloS one 10, e0126980, doi:10.1371/journal.pone.0126980 (2015).

64. Doring, G., Flume, P., Heijerman, H. & Elborn, J. S. Treatment of lung infection in patients with cystic fibrosis: current and future strategies. Journal of cystic fibrosis : official journal of the European Cystic Fibrosis Society 11, 461–479, doi:10.1016/j.jcf.2012.10.004 (2012).

65. Cipolla, D., Blanchard, J. & Gonda, I. Development of Liposomal Ciprofloxacin to Treat Lung Infections. Pharmaceutics 8, doi:10.3390/pharmaceutics8010006 (2016).

66. Kuti, J. L., Nightingale, C. H., Knauft, R. F. & Nicolau, D. P. Pharmacokinetic properties and stability of continuous-infusion meropenem in adults with cystic fibrosis. Clinical therapeutics 26, 493–501, doi:10.1016/s0149-2918(04)90051-3 (2004).

67. Moriarty, T. F., McElnay, J. C., Elborn, J. S. & Tunney, M. M. Sputum antibiotic concentrations: implications for treatment of cystic fibrosis lung infection. Pediatric pulmonology 42, 1008–1017, doi:10.1002/ppul.20671 (2007).

68. De Baets, F., Schelstraete, P., Van Daele, S., Haerynck, F. & Vaneechoutte, M. *Achromobacter xylosoxidans* in cystic fibrosis: prevalence and clinical relevance. Journal of cystic fibrosis : official journal of the European Cystic Fibrosis Society 6, 75–78, doi:10.1016/j.jcf.2006.05.011 (2007).

69. Edwards, B. D. et al. Prevalence and Outcomes of *Achromobacter* Species Infections in Adults with Cystic Fibrosis: a North American Cohort Study. Journal of clinical microbiology 55, 2074–2085, doi:10.1128/jcm.02556-16 (2017).

70. Somayaji, R. et al. Clinical Outcomes Associated with *Burkholderia cepacia* Complex Infection in Patients with Cystic Fibrosis. Annals of the American Thoracic Society 17, 1542–1548, doi:10.1513/AnnalsATS.202003-204OC (2020).

71. Aspenberg, M., Sasane, S. M., Nilsson, F., Brown, S. P. & Waldetoft, K. W. Hygiene Hampers Competitive Release of Resistant Bacteria in the Commensal Microbiota. bioRxiv (2019).

72. Wale, N. et al. Resource limitation prevents the emergence of drug resistance by intensifying within-host competition. Proceedings of the National Academy of Sciences of the United States of America 114, 13774–13779, doi:10.1073/pnas.1715874115 (2017).

73. de Roode, J. C., Culleton, R., Bell, A. S. & Read, A. F. Competitive release of drug resistance following drug treatment of mixed *Plasmodium chabaudi* infections. Malaria journal 3, 33, doi:10.1186/1475-2875-3-33 (2004).

74. Bray, J. R. & Curtis, J. T. An Ordination of the Upland Forest Communities of Southern Wisconsin. Ecological Monographs 27, 325–349, doi:doi.org/10.2307/1942268 (1957).

75. Clarke, K. R. Non-parametric multivariate analyses of changes in community structure. Australian Journal of Ecology 18, 117–143, doi:doi.org/10.1111/j.1442-9993.1993.tb00438.x (1993).

76. Oksanen, J. et al. (2019).

77. van der Loo, M. P. The stringdist package for approximate string matching. https://CRAN.R-project.org/package=stringdist. . The R Journal 6, 111–122 (2014).

78. European Committee for Antimicrobial Susceptibility Testing of the European Society of Clinical, M. & Infectious, D. Determination of minimum inhibitory concentrations (MICs) of antibacterial agents by broth dilution. Clinical Microbiology and Infection 9, ix–xv, doi:doi.org/10.1046/j.1469-0691.2003.00790.x (2003).

79. McKenney, E. S. et al. Lipophilic Prodrugs of FR900098 Are Antimicrobial against *Francisella novicida In Vivo* and *In Vitro* and Show GlpT Independent Efficacy. PloS one 7, e38167, doi:10.1371/journal.pone.0038167 (2012).

80. Estrela, S. & Brown, S. P. Community interactions and spatial structure shape selection on antibiotic resistant lineages. PLoS computational biology 14, e1006179, doi:10.1371/journal.pcbi.1006179 (2018).

81. Brook, I. Beta-lactamase-producing bacteria in mixed infections. Clinical microbiology and infection : the official publication of the European Society of Clinical Microbiology and Infectious Diseases 10, 777–784, doi:10.1111/j.1198-743X.2004.00962.x (2004).

82. Brook, I., Pazzaglia, G., Coolbaugh, J. C. & Walker, R. I. *In-vivo* protection of group A beta-haemolytic streptococci from penicillin by beta-lactamase-producing *Bacteroides* species. The Journal of antimicrobial chemotherapy 12, 599–606, doi:10.1093/jac/12.6.599 (1983).

83. Dugatkin, L. A., Perlin, M., Lucas, J. S. & Atlas, R. Group-beneficial traits, frequency-dependent selection and genotypic diversity: an antibiotic resistance paradigm. *Proceedings*. Biological sciences 272, 79–83, doi:10.1098/rspb.2004.2916 (2005).

84. Hackman, A. S. & Wilkins, T. D. *In vivo* protection of *Fusobacterium necrophorum* from penicillin by *Bacteroides fragilis*. Antimicrobial agents and chemotherapy 7, 698–703, doi:10.1128/aac.7.5.698 (1975).

85. Tacking, R. Penicillinase-producing bacteria in mixed infections in rabbits treated with penicillin. 2. Studies on penicillinase-producing coli- and pyocyaneus-bacteria. Acta pathologica et microbiologica Scandinavica 35, 445–454 (1954).

86. May, R. M. Thresholds and breakpoints in ecosystems with a multiplicity of stable states. Nature 269, 471–477, doi:10.1038/269471a0 (1977).

87. Vandeplassche, E., Tavernier, S., Coenye, T. & Crabbé, A. Influence of the lung microbiome on antibiotic susceptibility of cystic fibrosis pathogens. European Respiratory Review 28, 190041, doi:10.1183/16000617.0041-2019 (2019).

88. Callahan, B. J. et al. DADA2: High-resolution sample inference from Illumina amplicon data. Nature methods, doi:10.1038/nmeth.3869 (2016).

89. Holden, M. T. et al. The genome of *Burkholderia cenocepacia* J2315, an epidemic pathogen of cystic fibrosis patients. Journal of bacteriology 191, 261–277, doi:JB.01230-08 [pii] 10.1128/JB.01230-08 [doi] (2009).

90. Bertness, M. D. & Callaway, R. Positive interactions in communities. Trends in ecology & evolution 9, 191–193, doi:10.1016/0169-5347(94)90088-4 (1994).

91. Hesse, E. et al. Stress causes interspecific facilitation within a compost community. Ecology letters, doi:10.1111/ele.13847 (2021).

92. Piccardi, P., Vessman, B. & Mitri, S. Toxicity drives facilitation between 4 bacterial species. Proceedings of the National Academy of Sciences of the United States of America 116, 15979–15984, doi:10.1073/pnas.1906172116 (2019).

93. Brown, M. R., Collier, P. J. & Gilbert, P. Influence of growth rate on susceptibility to antimicrobial agents: modification of the cell envelope and batch and continuous culture studies. Antimicrobial agents and chemotherapy 34, 1623–1628, doi:10.1128/aac.34.9.1623 (1990).

94. Gilbert, P., Collier, P. J. & Brown, M. R. Influence of growth rate on susceptibility to antimicrobial agents: biofilms, cell cycle, dormancy, and stringent response. Antimicrobial agents and chemotherapy 34, 1865–1868, doi:10.1128/aac.34.10.1865 (1990).

95. Baumgartner, M., Bayer, F., Pfrunder-Cardozo, K. R., Buckling, A. & Hall, A. R. Resident microbial communities inhibit growth and antibiotic-resistance evolution of *Escherichia coli* in human gut microbiome samples. Plos Biol 18, e3000465, doi:10.1371/journal.pbio.3000465 (2020).

96. Wu, Y., Klapper, I. & Stewart, P. S. Hypoxia arising from concerted oxygen consumption by neutrophils and microorganisms in biofilms. Pathogens and disease 76, doi:10.1093/femspd/fty043 (2018).

97. Cornforth, D. M., Diggle, F. L., Melvin, J. A., Bomberger, J. M. & Whiteley, M. Quantitative Framework for Model Evaluation in Microbiology Research Using *Pseudomonas aeruginosa* and Cystic Fibrosis Infection as a Test Case. mBio 11, doi:10.1128/mBio.03042-19 (2020).

98. Flynn, J. M. et al. Disruption of Cross-Feeding Inhibits Pathogen Growth in the Sputa of Patients with Cystic Fibrosis. mSphere 5, doi:10.1128/mSphere.00343-20 (2020).

99. Quinn, R. A. et al. Niche partitioning of a pathogenic microbiome driven by chemical gradients. Science advances 4, eaau1908, doi:10.1126/sciadv.aau1908 (2018).

100. Banerjee, S., Schlaeppi, K. & van der Heijden, M. G. A. Keystone taxa as drivers of microbiome structure and functioning. Nat Rev Microbiol 16, 567–576, doi:10.1038/s41579-018-0024-1 (2018).

101. Allen, R. C., McNally, L., Popat, R. & Brown, S. P. Quorum sensing protects bacterial co-operation from exploitation by cheats. The ISME journal 10, 1706–1716, doi:10.1038/ismej.2015.232 (2016).

102. Pollitt, E. J., West, S. A., Crusz, S. A., Burton-Chellew, M. N. & Diggle, S. P. Cooperation, quorum sensing, and evolution of virulence in *Staphylococcus aureus*. Infect Immun 82, 1045–1051, doi:10.1128/iai.01216-13 (2014).

103. Limoli, D. H. et al. *Staphylococcus aureus* and *Pseudomonas aeruginosa* co-infection is associated with cystic fibrosis-related diabetes and poor clinical outcomes. European journal of clinical microbiology & infectious diseases : official publication of the European Society of Clinical Microbiology 35, 947–953, doi:10.1007/s10096-016-2621-0 (2016).

104. Maliniak, M. L., Stecenko, A. A. & McCarty, N. A. A longitudinal analysis of chronic MRSA and *Pseudomonas aeruginosa* co-infection in cystic fibrosis: A single-center study. Journal of cystic fibrosis : official journal of the European Cystic Fibrosis Society 15, 350–356, doi:10.1016/j.jcf.2015.10.014 (2016).

105. Leibold, M. A. et al. The metacommunity concept: a framework for multi-scale community ecology. Ecology letters 7, 601–613 (2004).

106. Wright, E. S. & Vetsigian, K. H. Inhibitory interactions promote frequent bistability among competing bacteria. Nature communications 7, 11274, doi:10.1038/ncomms11274 (2016).

107. Read, A. F., Day, T. & Huijben, S. The evolution of drug resistance and the curious orthodoxy of aggressive chemotherapy. Proceedings of the National Academy of Sciences of the United States of America 108 **Suppl 2**, 10871–10877, doi:10.1073/pnas.1100299108 (2011).

108. Halpin, A. L. et al. Intestinal microbiome disruption in patients in a long-term acute care hospital: A case for development of microbiome disruption indices to improve infection prevention. American journal of infection control 44, 830–836, doi:10.1016/j.ajic.2016.01.003 (2016).

109. McAdams, D., Wollein Waldetoft, K., Tedijanto, C., Lipsitch, M. & Brown, S. P. Resistance diagnostics as a public health tool to combat antibiotic resistance: A model-based evaluation. Plos Biol 17, e3000250, doi:10.1371/journal.pbio.3000250 (2019).

110. Moran Losada, P. et al. The cystic fibrosis lower airways microbial metagenome. ERJ open research 2, doi:10.1183/23120541.00096-2015 (2016).

111. Mitchell, J. *Streptococcus mitis*: walking the line between commensalism and pathogenesis. Molecular oral microbiology 26, 89–98, doi:10.1111/j.2041-1014.2010.00601.x (2011).

112. Mathee, K. et al. Mucoid conversion of *Pseudomonas aeruginosa* by hydrogen peroxide: a mechanism for virulence activation in the cystic fibrosis lung. *Microbiology (Reading*, England) 145 **( Pt** **6**), 1349–1357 (1999).

113. Martin, D. W. et al. Mechanism of conversion to mucoidy in *Pseudomonas aeruginosa* infecting cystic fibrosis patients. Proceedings of the National Academy of Sciences of the United States of America 90, 8377–8381, doi:10.1073/pnas.90.18.8377 (1993).

114. Ruddy, J. et al. Sputum tobramycin concentrations in cystic fibrosis patients with repeated administration of inhaled tobramycin. Journal of aerosol medicine and pulmonary drug delivery 26, 69–75, doi:10.1089/jamp.2011.0942 (2013).

115. Wenzler, E. et al. Meropenem-RPX7009 Concentrations in Plasma, Epithelial Lining Fluid, and Alveolar Macrophages of Healthy Adult Subjects. Antimicrobial agents and chemotherapy 59, 7232–7239, doi:10.1128/aac.01713-15 (2015).

116. Caporaso, J. G. et al. Global patterns of 16S rRNA diversity at a depth of millions of sequences per sample. Proceedings of the National Academy of Sciences of the United States of America 108 **Suppl 1**, 4516–4522, doi:10.1073/pnas.1000080107 (2011).

117. Stoddard, S. F., Smith, B. J., Hein, R., Roller, B. R. & Schmidt, T. M. rrnDB: improved tools for interpreting rRNA gene abundance in bacteria and archaea and a new foundation for future development. Nucleic Acids Res 43, D593–598, doi:10.1093/nar/gku1201 (2015).

118. R: a language and environment for statistical computing. (R Foundation for Statistical Computing 2018).

119. Wickham, H. ggplot2-Elegant Graphics for Data Analysis. Springer International Publishing. *Cham*, Switzerland (2016).

120. Van den Boogaart, K., Tolosana, R. & Bren, M. compositions: Compositional Data Analysis. R package version 2.0-0. R Foundation for Statistical Computing., Vienna (2020).

121. Kassambara, A. & Mundt, F. (R Foundation for Statistical Computing. Vienna, 2020).

